# Evolution of mechanisms controlling epithelial morphogenesis across animals: new insights from dissociation - reaggregation experiments in the sponge *Oscarella lobularis*

**DOI:** 10.1101/2021.03.22.436370

**Authors:** Amélie Vernale, Maria Mandela Prünster, Fabio Marchianò, Henry Debost, Nicolas Brouilly, Caroline Rocher, Dominique Massey-Harroche, Emmanuelle Renard, André Le Bivic, Bianca H. Habermann, Carole Borchiellini

## Abstract

**Background:** The ancestral presence of epithelia in Metazoa is no longer debated. Even though Porifera seem to be the best candidates to be the sister group to all other Metazoa, hardly anything is known about the proteins involved in the composition of cell-cell junctions or about the mechanisms that regulate epithelial morphogenetic processes in this phylum.

**Results:** To get insights into the early evolution of epithelial morphogenesis, we focused on morphogenic characteristics of the homoscleromorph sponge *Oscarella lobularis.* Homoscleromorpha are a sponge class with a typical basement membrane and adherens-like junctions unknown in other sponge classes. We took advantage of the dynamic context provided by cell dissociation-reaggregation experiments to explore morphogenetic processes in epithelial cells in an early lineage by combining fluorescent and electronic microscopy observations and RNA sequencing approaches at key time-points of the dissociation and reaggregation processes.

**Conclusions:** Our results show that part of the molecular toolkit involved in the loss and restoration of epithelial features such as cell-cell and cell-matrix adhesion is conserved between Homoscleromorpha and Bilateria, suggesting their common role in the last common ancestor of animals. In addition, Sponge-specific genes are differently expressed during the dissociation and reaggregation processes, calling for future functional characterization of these genes.

## Background

The presence of epithelial tissues is a fundamental feature of all Metazoa. They are the first a visible sign of cellular differentiation as early as the blastula stage and a starting point for building animal bodies during development. The emergence of epithelia, over 800 million years ago, represent a pivotal evolutionary innovation at the origin of the establishment of a permanent multicellularity in the last metazoan common ancestor (LMCA) (Bich et al., 2019; Brunet and King, 2017; Leys and Riesgo, 2012; Leys et al., 2009; Miller et al., 2013; Nielsen, 2012; Tyler, 2003; Renard et al., in press).

According to bilaterian features, an epithelium is usually defined as a layer of cells i) showing coordinated polarity controlled by three polarity complexes (Crumbs, Scribble and Par); ii) connected by cell-cell junctions (adhesive, communicating and sealing junctions) and iii) attached by their basal pole to a basement membrane (containing type IV Collagen) via cell-matrix junctions involving Integrins (Tyler, 2003) (Figure 1A). Such a typical bilaterian organization of the epithelium, fitting all three mentioned criteria, is however not commonly present in non-bilaterian lineages except in Cnidaria (reviewed in Renard et al., in press) (Figure 1B). It is therefore necessary to characterize and compare epithelial features of the three remaining non-bilaterian phyla (Porifera, Ctenophora and Placozoa). This will allow to understand how and when the cnidarian-bilaterian epithelial features emerged.

**Figure 1:**
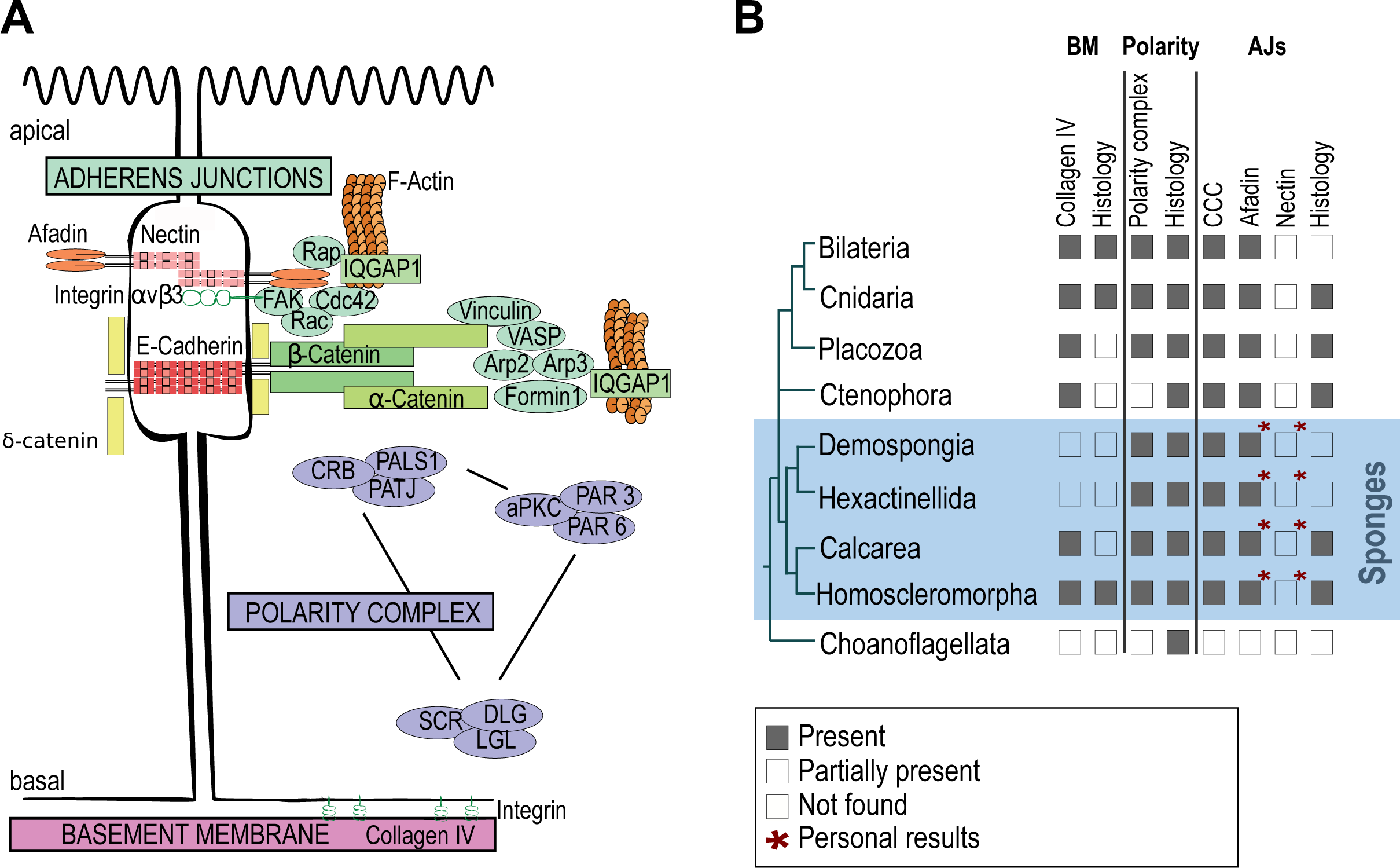
The main molecular actors of epithelial cell features in bilaterians and their presence/absence in non-bilaterian metazoans. **(A)** Schematic representation (modified from (Miyoshi and Takai, 2011) of typical bilaterian epithelial cells and of proteins involved in (i) adherens junctions, (ii) the three polarity complexes and (iii) the basement membrane (Review in Renard et al., in press). (i) In detail, cell adhesion can be Nectin- and E-cadherin-based and connect via numerous proteins to the actin cytoskeleton: Nectins via Afadin and Integrin; E-adherins via δ-, β- and α-catenins to the cell adhesion complex proteins Vinculin, VASP, Formin, Arp2/3. In addition, Rap, Rac, CDC42, FAK and IqGAP1 regulate the changes and organization of the actin skeleton (Miyoshi and Takai, 2011). (ii) The three major polarity complexes namely the apically located PAR3/PAR6/aPKC complex and the CRUMBS (CRB)/PALS1/PATJ complex and along with a lateral SCRIBBLE (SCR)/DLG/LGL complex (Assémat et al., 2008). iii) The basement membrane of bilaterians involves mainly type IV collagen, laminins, perlecan and nidogen; laminins interact with integrins to establish cell-matrix adhesion (Fidler et al., 2017). The nomenclature chosen in the scheme is according to human proteins. **(B**) Presence/absence of epithelial histological features and epithelial genes in non-bilaterians mapped on a schematic representation of consensual phylogenetic relationships among metazoans according to the literature (for review see Kapli and Telford, 2020; Schenkelaars et al., 2019; Renard et al., in press). “Partially present” means that either only part of the genes were found in the considered taxa or that some species of the taxa lack the genes or features.

Albeit phylogenetic relationships at the basis of the animal tree are being thoroughly discussed for more than ten years, Porifera (sponges) are probably the most basal metazoan lineage (Daley and Antcliffe, 2019; Erives and Fritzsch, 2019; Feuda et al., 2017; Francis and Canfield, 2020; Kapli and Telford, 2020; King and Rokas, 2017; Littlewood, 2017; Schenkelaars et al., 2019; Simion et al., 2017). Porifera therefore constitute a pivotal group to trace the emergence of epithelia.

The phylum Porifera is divided into 4 distinct classes (Calcarea, Demospongiae, Hexactinellida and Homoscleromorpha) harboring distinct epithelial features according to histological observations (Ereskovsky, 2010; Ereskovsky et al., 2009; Fidler et al., 2017; Leys and Riesgo, 2012; Leys et al., 2009) (Figure 1B). Among them, only homoscleromorph sponges (Homoscleromorpha) possess tissues that fully meet the previous definition of an epithelium. Indeed, the 3 other sponge classes (Demospongiae, Hexactinellida and Calcarea) are devoid of a basement membrane and adherens-like junctions (Ereskovsky et al., 2009; Leys and Riesgo, 2012; Leys et al., 2009; Nielsen, 2008; Tyler, 2003).

Despite these histological differences, comparative genomic and transcriptomic analyses evidenced that all 4 sponge classes possess a whole set of genes encoding core proteins needed to build bilaterian epithelia (catenins, cadherins, polarity complexes) (Belahbib et al., 2018; Fahey and Degnan, 2010; Francis et al., 2017; Kenny et al., 2020; Nichols et al., 2012; Riesgo et al., 2014). According to this discrepancy between histological and genetic data (reviewed in Renard et al., 2018) and despite recent efforts to solve this issue by modern immunodetection and proteomic approaches (Miller et al., 2018; Mitchell and Nichols, 2019; Schippers et al., 2018), it is still unknown which proteins compose sponge cell-cell junctions and which molecular mechanisms are involved in epithelial morphogenetic processes in sponges (Green et al., 2020). Moreover, experimental and functional approaches are currently lacking to establish links between candidate genes of epithelial function and their effective purpose in the cell.

To explore the molecular mechanisms involved in epithelial morphogenesis, we focused on the homoscleromorph sponge *Oscarella lobularis* and we took advantage of the dynamic context offered by cell dissociation-reaggregation experiments (Curtis, 1962; Galtsoff, 1925; Humphreys, 1963; Huxley, 1921; Leith, 1979; Maclennan and Dodd, 1967; Rocher et al., 2020; Wilson, 1907). We carried out cell dissociation-reaggregation experiments at in buds (resulting from asexual reproduction), taking advantage of the high number of replicates and the transparency of tissue enabling fluorescent imaging (Rocher et al., 2020). The observation of the dissociation and the reaggregation processes at different time-points (by confocal imaging and electronic microscopy) enabled us to monitor the time-scale of the processes and determine time-points of key events (Figure 2). We focused on the disassembly and restoration of the internal epithelium, the choanoderm. This tissue is composed of choanocytes, a highly specialized cell-type organized in hollow spheres named choanocyte chambers. This tissue displays epithelial properties: the presence of a basement membrane composed of type IV Collagen and highly polarized cells establishing cell-cell junctions histologically similar to adherens junctions (Boury-Esnault and Rützler, 1997; Boury-Esnault et al., 2003; Boute et al., 1996; Ereskovsky, 2010; Ereskovsky and Boury-Esnault, 2002; Ereskovsky et al., 2009; Gazave et al., 2012; Leys and Riesgo, 2012; Leys et al., 2009; Rocher et al., 2020). In parallel, we performed high-throughput RNA sequencing at selected time-points (Figure 2) and looked at temporal expression dynamics to identify genes involved in epithelial patterning (loss of adhesion and polarity, and re-epithelization) using a *de novo* assembled and annotated *Oscarella lobularis* transcriptome. This unprecedent approach to study epithelial morphogenesis in a sponge provides a large set of new data and insights on the evolution of the metazoan toolkit involved in epithelial morphogenesis.

**Figure 2:**
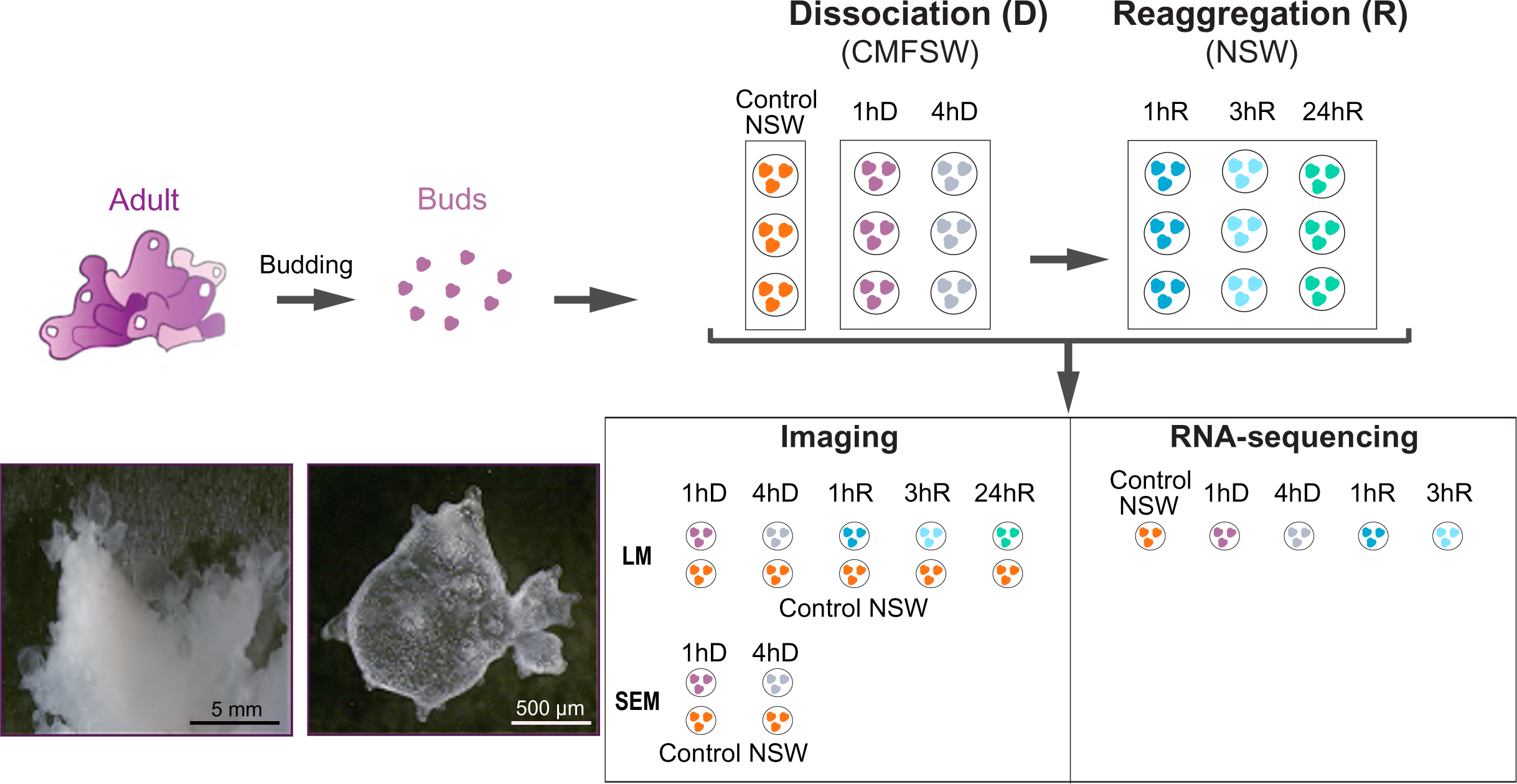
Experimental strategy for the time course study of cell-dissociation and reaggregation processes in the homoscleromorph sponge *Oscarella lobularis.* Adults of *Oscarella lobularis* can reproduce asexually by budding, forming so-called buds. The bud tissues are highly similar to the adult tissues and buds were shown to be more convenient for experiments (Ereskovsky and Tokina, 2007; Rocher et al., 2020). Clonal buds from one adult were put in Calcium-Magnesium-Free Sea Water (CMFSW) to initiate cell dissociation. After 4 hours of dissociation buds were placed back into Natural Sea Water (NSW) to initiate reaggregation. Samples were collected at different times for imaging (Scanning Electron Microscopy (SEM), Light confocal Microscopy (LM)) and RNA-sequencing. Abbreviated as follows: 1hour (1hD) and 4hours (4hD) of dissociation and 1hour (1hR), 3 hours (3hR) and 24 hours (24hR) of reaggregation.

## Results

### Loss of epithelial features in the choanoderm during dissociation

Live buds were first incubated with fluorescent labeled lectin PhaE (*Phaseolus vulgaris Erythroagglutinin*) to specifically stain choanocytes (Borchiellini et al., 2021; Rocher et al., 2020) (Supplementary Figure S1). Then, cell dissociation was induced by depletion of divalent cations in calcium-magnesium-free artificial sea water (CMFSW). As a control, we used buds incubated in natural sea water (NSW) in culture plates for the same duration. Choanoderm features were observed by both, confocal microscopy imaging (after type IV Collagen or acetylated tubulin immunostaining, phalloidin and DAPI counter-staining) and Scanning Electron Microscopy (SEM-3D).

Whatever the incubation times of the control (NSW), the choanocyte chambers looked like as described before in this species (Boury-Esnault et al., 2003; Ereskovsky and Boury-Esnault, 2002; Rocher et al., 2020): they appeared as a rounded and organized structures (Figure 3A, C). Choanocytes had a canonical shape and were highly polarized. Their larger basal side rested on a basal lamina and they had an actin-rich collar microvilli surrounding a flagellum at their thinner apical side (Figure 3A, C, C’, C’’). Cell-cell contacts could be identified as an actin-rich ring around each cell (Figure 3B). 3D reconstructions of SEM images showed that cell-cell contacts on the lateral side were almost inexistent below the actin ring, indicating that the tissue cohesion mainly relied on these thin lateral adherens-like junctions and on a large area of latero-basal cell-matrix contacts (Figure 3C’, C’’).

**Figure 3:**
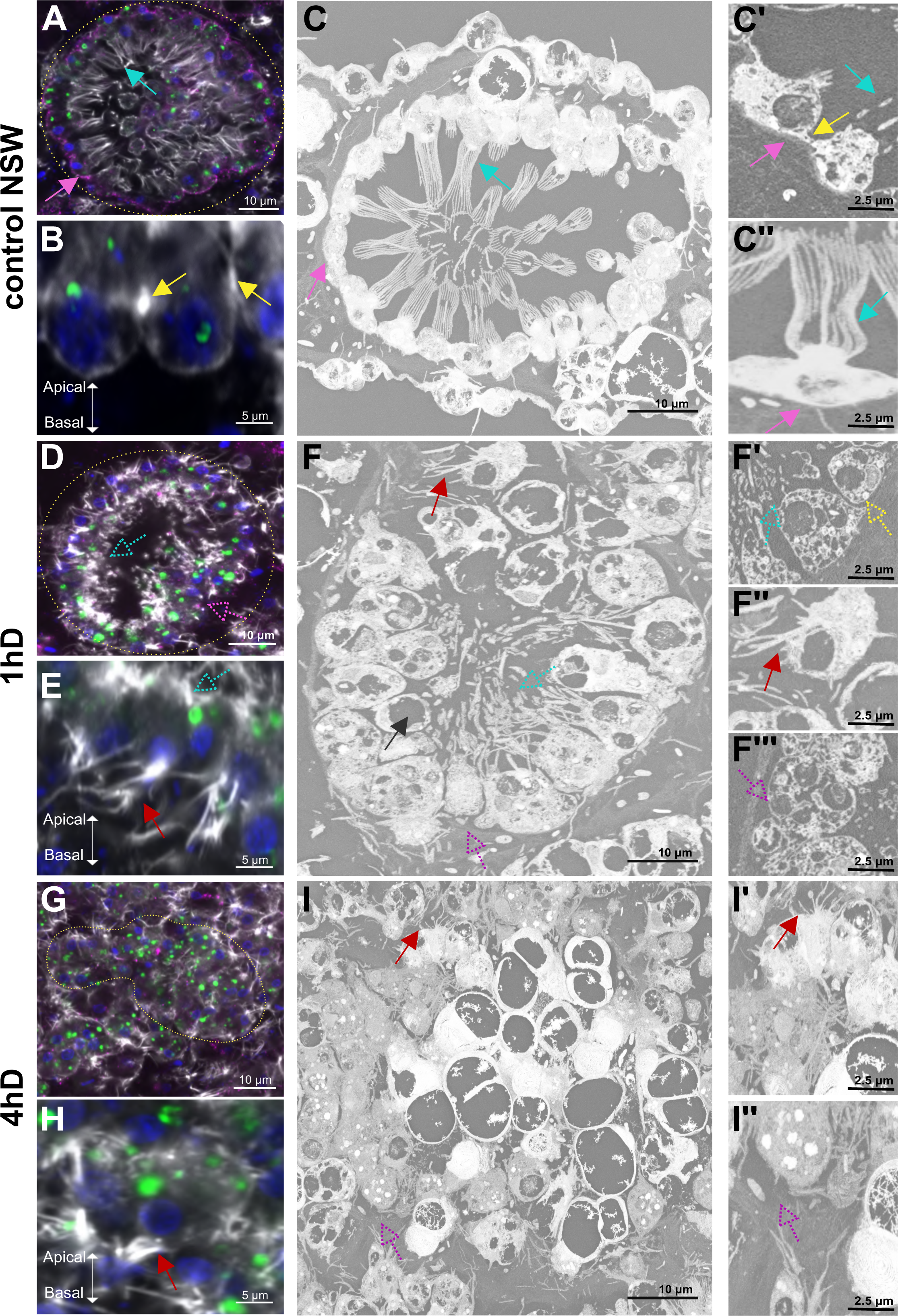
Structural changes in the choanoderm during dissociation. **(A- C’’)** Control conditions in natural sea water (1h NSW): **(A)** Confocal microscopic view (LM) of a choanocyte chamber, the classical round shape of the choanocyte chamber is visible (dotted yellow line) with its typical collar of apical microvilli (blue arrow). The type IV Collagen staining (in magenta) is lining the basal pole of choanocyte chambers (pink arrow). **(A)** LM focused view on the actin-rich cellular junction between choanocytes (yellow arrow). **(B)** Scanning Electron Microscopic (SEM) view of a choanocyte chamber with collar of apical microvilli (cyan arrow), and a basement membrane lining the choanocyte chamber (pink arrow). **(C’)** Focus on the cell-cell junction (yellow arrow), basal lamina (pink arrow) and apical microvilli (cyan arrow). **(C’’)** Choanocyte with its complete apical microvilli collar (cyan arrow) and complete basal lamina (pink arrow). **(D-F’’’)** Choanocyte chambers observed one hour after incubation in calcium-magnesium-free sea water (1hD CMSFW): **(D)** The general architecture of the choanocyte chamber is still observable in LM (dotted yellow line), immunostaining of type IV Collagen is not visible anymore (dotted pink arrow), apical microvilli are disintegrated (dotted cyan arrow). **(E)** Focus (LM) on a choanocyte with its disintegrated apical microvilli (dotted cyan arrow) and numerous basal actin protrusions (red arrow). **(F)** SEM view of a choanocyte chamber. Choanocytes present vacuoles (black arrow), disintegrated apical microvilli (dotted cyan arrow) and protrusions (red arrow). The basal lamina is detached from the choanocytes (dotted pink arrow). **(F’)** Focus (SEM) on the disintegration of the apical microvilli (cyan dotted arrow) and on the loss of cell-cell contact (dotted yellow arrow). **(F’’)** Focus on basal actin protrusions at the basal pole of a choanocyte (SEM) (red arrow). **(F’’’)** Focus on two choanocytes with a clear detachment from the basal lamina (dotted pink arrow). **(G-I’’)** Choanocyte chambers observed four hours after incubation in calcium-magnesium-free sea water (4hD CMFSW): **(G)** LM view of a destructured choanocyte chamber, immunostaining of type IV Collagen is not visible and the general architecture of the choanocyte chamber is no more recognizable (yellow dotted line). **(H)** (LM) Focus on a choanocyte with numerous actin basal protrusions (red arrow). **(I**) SEM view of choanocyte showing actin protrusions (red arrow) and the whole destructuring of the basal lamina (dotted pink arrow). **(I’)** (SEM) focus on the actin protrusions (red arrow). **(I’’)** The basal lamina is completely lost at this time-point (SEM) (dotted pink arrow). For all confocal (LM) pictures: Staining in grey: Phalloidin, Green: PhaE lectin, Blue: DAPI, Magenta: type IV Collagen. Additional picture of control condition in NSW is available in the Supplementary Figure S2B.

After incubating the buds for either 20 minutes, 1, 2, 3 or 4 hours in CMFSW, we identified two main steps in the dissociation process:

- After 1 hour of incubation in CMFSW (1hD), the choanocyte chambers could still be identified and were forming cohesive structures (Figure 3D, F). However, the shape of choanocytes was strongly affected compared to the control condition: they adopted a round shape and the collar of actin microvilli at their apical pole lost its organization. The microvilli displayed some degree of fragmentation (Figure 3D, E, F, F’; Supplementary Figure S2A-1hD). At this stage, 99.4% of the choanocyte chambers were affected (n= 178; Supplementary Table S1A). In addition, large vacuoles were present in choanocytes (Figure 3F). In parallel, F-actin containing protrusions were much more numerous at the basal pole of choanocytes (Figure 3E, F, F’’) and cell adhesion, including both, cell-cell contacts (Figure 3F’) as well as cell-basal lamina contacts, (Figure 3F’’’) were lost. This disorganization of the basement membrane was also evidenced by the decrease of type IV Collagen labeling (Figure 3D) surrounding these structures.
- After 4 hours of calcium/magnesium deprivation (4hD), the choanocyte chambers were disorganized and only remained recognizable due to PhaE staining. They formed compact irregular pseudo-spheres, the internal cavity and the former monolayered epithelial organization of the choanoderm were no longer present in any of the observed chambers (Figure 3G, I; Supplementary figure S2A-4hD and B-4hD for the controls). At this stage, 100% of the choanocyte chambers were unstructured (n= 216; Supplementary Table S1B). The apical collar of microvilli was completely absent (Figure 3G, H). F-actin protrusions were still present at the basal pole of these cells (Figures 3H, I, I’). The flagellum (anti acetylated tubulin immunostaining) was still present despite the absence of a clear polarized cell-body (Supplementary Figure S3A). Moreover, type IV Collagen staining was lost (Figure 3G) and the basement membrane was not visible anymore (Figure 3I, I”).

All phenotypes observed were reproducible between different individuals as well as between the majority of choanocyte chambers of the same bud (see images at lower magnification provided in Supplementary Figure S2A; Statistical analyses provided in Supplementary Table S1).

In conclusion, after 4 hours of calcium-magnesium deprivation, the main epithelial features of the choanoderm were lost, defined by the loss of the basement membrane, of cell-cell junctions, of cell-matrix contacts and of a clear cell polarity.

### Reaggregation, reorganization and re-epithelialization of the choanoderm

Buds dissociated previously by incubation for 4 hours in CMFSW were put back into NSW. By monitoring the reaggregation process at different time-points (30 min, 1, 2, 3, 5, 6, 24 hours), we were able to define 3 main steps:

- After 1 hour incubation in NSW (1hR) (Supplementary Figure S2C; Control in S2B-1hR), PhaE-stained cells (previous choanocytes) started to realign, although the structure of the chamber was not recovered at this step compared to the control.
- After 3 hours in NSW (3hR) (Figure 4A, B; Control in Supplementary Figure S2B-3hR) an accumulation of F-actin detectable in the center of PhaE-stained cells indicated not only the reformation of a small central cavity but also the reformation of a microvillar collar: cell polarity was already re-established at that time-point and the cells began to reorganize into an monostratified cell layer (Figure 4A, B). Choanocytes re-established contacts with each other and with the basement membrane that began to recover. A discontinuous, spot-like type IV Collagen immunostaining could again be observed around these structures. Thus, at that time-point the main epithelial features were being recovered: basement membrane, cell-matrix and cell-cell adhesion, as well as cell polarity. The cells were not only reaggregated and reorganized (compared to 1hR) but underwent a re-epithelialization process.
- After 24 hours in NSW (24hR), 96.8% of choanocyte chambers recovered their initial structure (n= 94, Supplementary Table S1C) and choanocytes regained their classical morphology (Figure 4C, D; Control in Supplementary Figure S2B-24hR). The basement membrane was recovered with a continuous type IV Collagen line surrounding the chambers, as well as a clear reformation of the apical microvilli.

**Figure 4:**
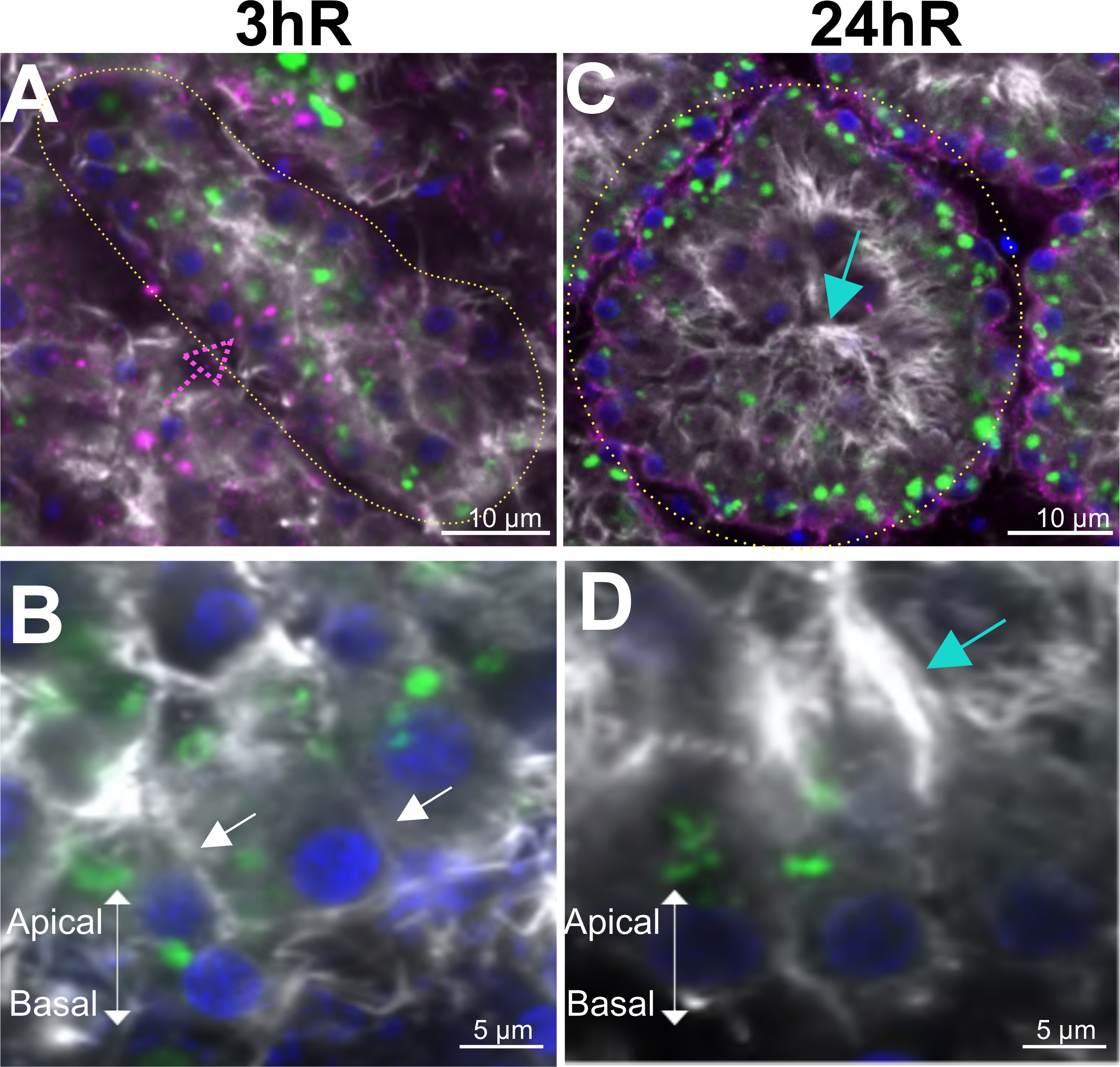
Restoration of the choanoderm during reaggregation. Confocal microscopic views of **(A)** a choanocyte chamber after 3 hours of reaggregation (3hR) in natural sea water (NSW), the immunostaining of type IV Collagen (in magenta) is discontinuous (dotted pink arrow). **(B)** Focus on three choanocytes starting to realign with each other at 3hR (white arrows). **(C)** A choanocyte chamber fully restructured after 24 hours of reaggregation (24hR), choanocytes present typical apical microvilli (cyan arrow). **(D)** Focus on choanocytes with a restored apical microvilli (cyan arrow). For all pictures, staining in grey: Phalloidin, in Green: Pha-E lectin, in Blue: DAPI, in Magenta: type IV Collagen. Additional views of control conditions in NSW are available in Supplementary Figures S2B.

In summary, the reaggregation of choanocytes resulted in a complete recovery of the normal morphology of the choanoderm, demonstrating the reversibility of the dissociation process.

### Characterization of the transcriptome of *Oscarella lobularis*

An earlier attempt to obtain the genome sequence of the *O. lobularis* genome resulted only in a partial, highly fragmented genome due to a high number of repetitive sequences and a large number of contaminant reads from bacteria and archaea (Belahbib, 2018). The estimated gene content of this partial resource was ∼17885 protein-coding genes, which is low compared to other sponges (Ereskovsky et al. 2017, Kenny et al., 2020). We therefore decided to sequence and *de novo* assemble the transcriptome of *O. lobularis,* attempting to obtain a near-complete transcriptome of this sponge species. To this end, two types of sequencing libraries were merged: one library was prepared from a reproductive adult with embryos and larvae previously sequenced by 454 sequencing (Schenkelaars et al., 2015); the second library was obtained from dissociating/reaggregating buds and from buds in NSW and sequenced by Illumina sequencing in the course of the present work. Sequencing of the two libraries resulted in 538 821 reads from 454 sequencing and 480 883 138 Illumina reads, respectively (Figure 5). Low quality reads with a Phred quality score < 20 were trimmed and removed from both libraries. All reads were submitted to the Short Read Archive at NCBI (Bioproject PRJNA659410).

**Figure 5:**
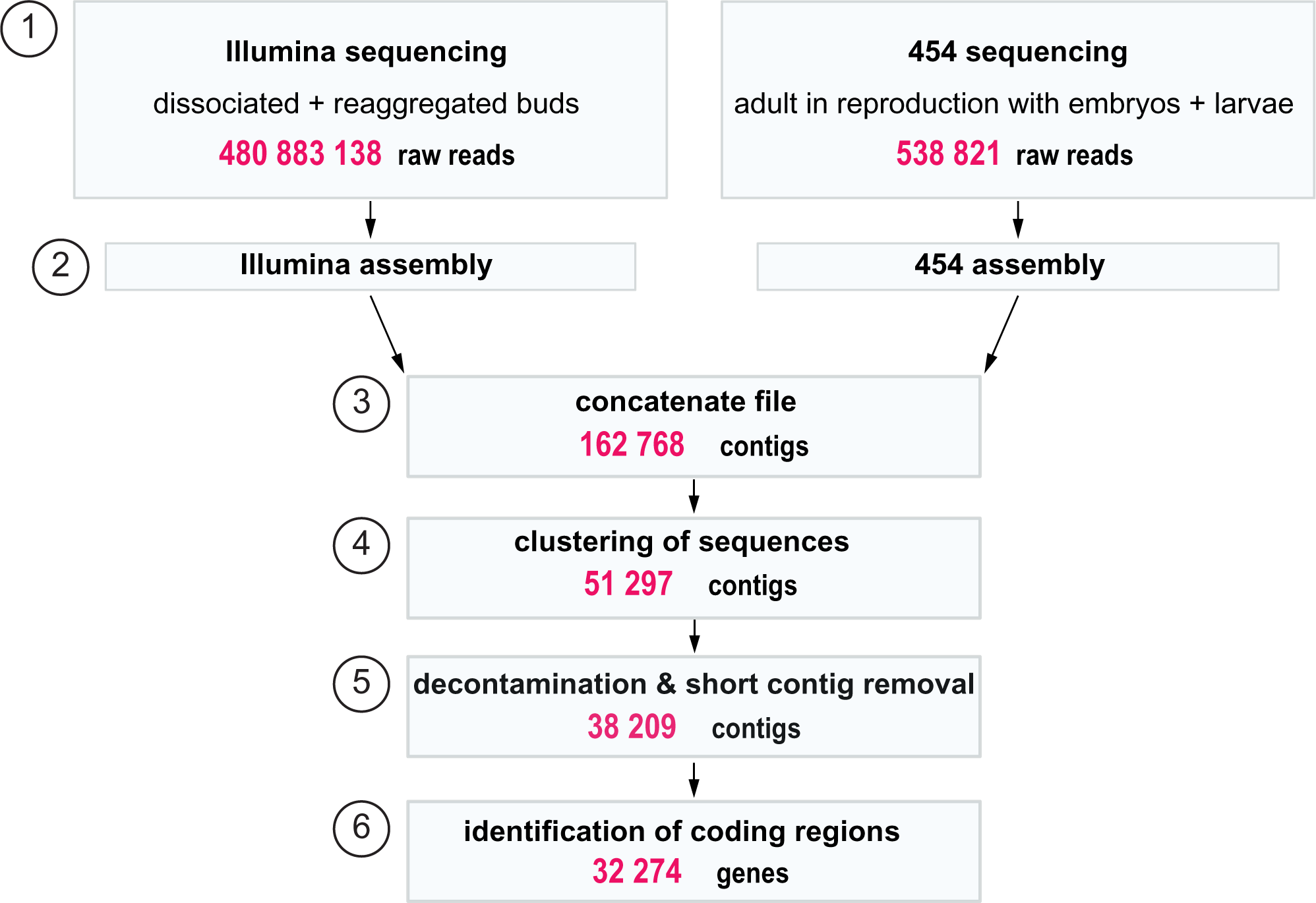
Schematic workflow of the steps performed for *de novo* transcriptome assembly of *Oscarella lobularis*. **(1)** Illumina sequencing was performed on biological samples of 40 pooled clonal stage 3 buds exposed to the same conditions: dissociated in CMFSW and reaggregated in NSW. 454 sequencing was performed on tissue from one adult undergoing sexual reproduction containing a mixed population of embryos and pre-larvae collected in the bay of Marseille (Schenkelaars et al., 2015). Illumina reads were quality controlled using FastQC (http://www.bioinformatics.babraham.ac.uk/projects/fastqc/). Low-quality reads and adapters were removed using TrimGalore (https://github.com/FelixKrueger/TrimGalore). Illumina and 454 sequencing libraries were first *de novo* assembled individually and then merged. lllumina sequences were *de novo* assembled using Trinity. 454 reads were *de novo* assembled with Staden and GAP4 (Bonfield et al., 1995) by Eurofins. **(2)** The two assemblies were then concatenated. **(3)** Redundancy was reduced by clustering contigs > 80% identity with CD-hit EST (Li and Godzik, 2006) retaining the longer contig. **(4)** To remove sequence redundancy further, we indexed and mapped the contigs with GMAP (Wu and Watanabe, 2005) with a 95% sequence identity. To remove contaminants, we used Vecscreen and BlastN. **(5)** Using Transdecoder (https://github.com/TransDecoder) with the Transdecoder.predict option we were able to identify the most likely protein sequences encoded in open reading frames (ORFs) ≥ 100 aa. For additional experimental details see the Material and Methods section.

*De novo* assembly of the two different libraries was done separately (see Material and Methods section and Figure 5), the concatenated dataset resulted in 162 768 contigs. After removing redundant contigs, as well as possible contaminations (Archaea, Bacteria, Viridiplantae, Fungi and Dinoflagellata) the final *O. lobularis* transcriptome contained 32 274 contigs. The average contig length was 958 bp, with the smallest contig having a length of 255 bp and the largest 47 628 bp. The mean GC content based on all contigs was 48.2% (Figure 6A).

**Figure 6:**
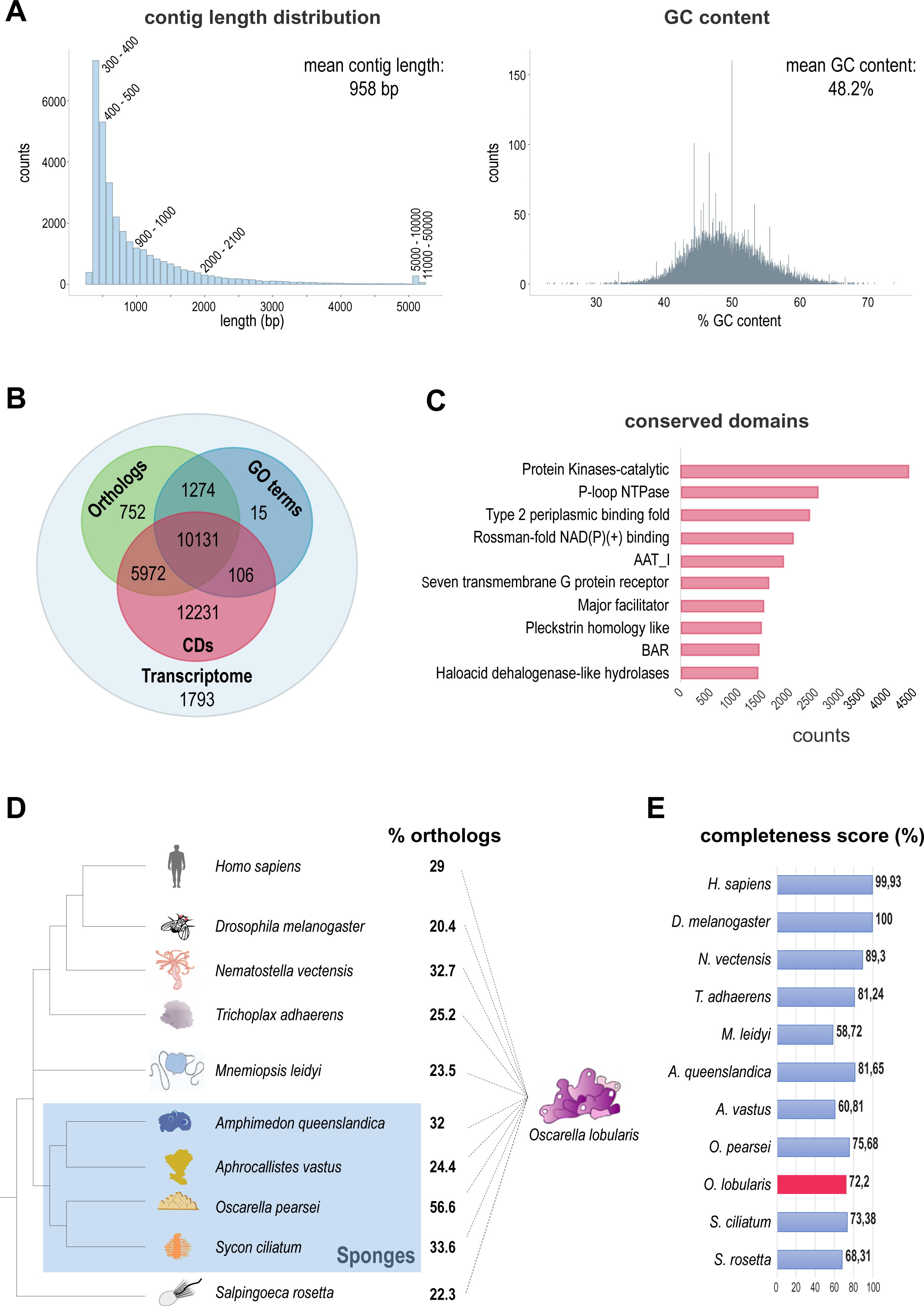
Characteristics of the *de novo* transcriptome assembly of *O. lobularis.* **(A)** Contig length distribution and GC content of the *O. lobularis* transcriptome from mixed stages and types of sequencing. Statistics was done after *de novo* assembly, concatenation, decontamination, and length cleaning steps. The smallest contig is 255 bp long and the longest contig has 47 628 bp coming from a single long ORF encoding a Mucin ortholog. The average contig length is 958 bp. The mean percentage of GC content is 48.2 %. **(B)** Annotated coding regions either with eggNOG (Huerta-Cepas et al., 2019) (orthologs) (18129 sequences), GO Terms (11526), and Conserved Domains (CDs) (Marchler-Bauer et al., 2011) (28440 sequences) (Supplementary Table S2). For 1793 contigs no described CD or ortholog could be found, corresponding to 5.5 % of the identified coding regions. **(C)** The ten most frequent Conserved Domains (CDs) in the *O. lobularis* transcriptome. Among the top ten CDs are kinases, nucleotide binding domains, domains that are involved in membrane transport or membrane binding as well as those inferring changes in cell shape. The protein kinase domain was most abundant and was present in 4057 coding regions. Occurrences were counted once per coding region. **(D)** Percentage of orthologs, *O. lobularis* has in common with other species. 29% orthologs are shared with human, 25.2% with *Trichoplax adhaerens.* Shared orthologs with other sponges vary from 24% to 57%. 56.6% of contigs from *O. lobularis* have a clear ortholog in its close relative, *O. pearsei* (Supplementary Table S3). **(E)** DOGMA analysis (Dohmen et al., 2016) to evaluate the completeness of different metazoan transcriptomes (Supplementary Table S5). For *O. lobularis*, we found 72.2% of expected CD arrangements. This is within the range of other sponge transcriptomes.

We functionally annotated the transcriptome using groups of orthologs, gene ontology terms (GO), as well as conserved domains (CDs). We found an orthologous group for 18 129 contigs (56%), could assign a GO term to 11 526 contigs (35.7%) and predicted a CD for 28 440 contigs (88.1%) (Figure 6B, Supplementary Table S2A-E). Protein kinases were the most abundant CD identified, occurring in 4057 contigs. Among the other top CDs were membrane-associated domains, domains involved in membrane transport or membrane binding as well as those inferring changes in cell shape (Figure 6C). We further investigated with Orthofinder (Emms and Kelly, 2019) the percentage of orthologs shared between *O. lobularis* transcripts from our assembly and several other metazoans (Figure 6D, Supplementary Table S3). Shared orthologs with other sponges ranged between 24.4 and 56.6%, whereby the highest percentage of orthologs was found for the closest relative of *O. lobularis*, *Oscarella pearsei*. Between 20 and 33% of orthologs were shared with non-sponge animal species, with 29% of orthologs found in *H. sapiens*. We estimated the completeness of this transcriptome to be rather high, compared to other sponge transcriptomes (Supplementary Table S4), as our transcriptome assembly had a BUSCO (Seppey et al., 2019) score of 82%. To further support the completeness of the *O. lobularis* transcriptome, we looked at the percentage of conserved domain arrangements using DOGMA (Dohmen et al., 2016) (Figure 6E). We found that 72.2% of expected CD arrangements were present in the *O. lobularis* transcriptome. This number is similar to the other sponge transcriptomes tested, which had a maximum of 81.65% expected CD arrangements found in *Amphimedon queenslandica* (Demospongiae) and a minimum of 60% found in *Aphrocallistes vastus* (Hexactinellida) (Supplementary Table S5).

To conclude, the *O. lobularis* transcriptome presented here shows a high degree of completeness, with most of the contigs being assigned to an orthologous group, identifying a conserved domain or a gene ontology term. Furthermore, BUSCO and DOGMA results support the comprehensiveness of our resource. The transcriptome is available in the Bioproject PRJNA659410 on NCBI.

### Identification of differentially expressed genes during the cell-dissociation-reaggregation process

To identify molecular players involved in epithelium organization, we performed time-series RNA-sequencing at selected key time-points of the dissociation-reaggregation process: 2 time-points during the dissociation, 1 hour (1hD) and 4 hours (4hD); and 2 time-points during the reaggregation / re-epithelialization process, 1 hour (1hR) and 3 hours (3hR) (Figure 2). After trimming of raw reads, we mapped the reads to the *de novo* assembled *O. lobularis* transcriptome using Kallisto (Bray et al., 2016) and Tximport (Soneson et al., 2015). Principal component analysis (PCA) of normalized read counts showed that the different time-point replicates cluster together, except for the first time-point of reaggregation (1hR), which was dispersed between 4hD and 3hR clusters, suggesting a variability in the process timing between pooled individuals (Figure 7A).

**Figure 7:**
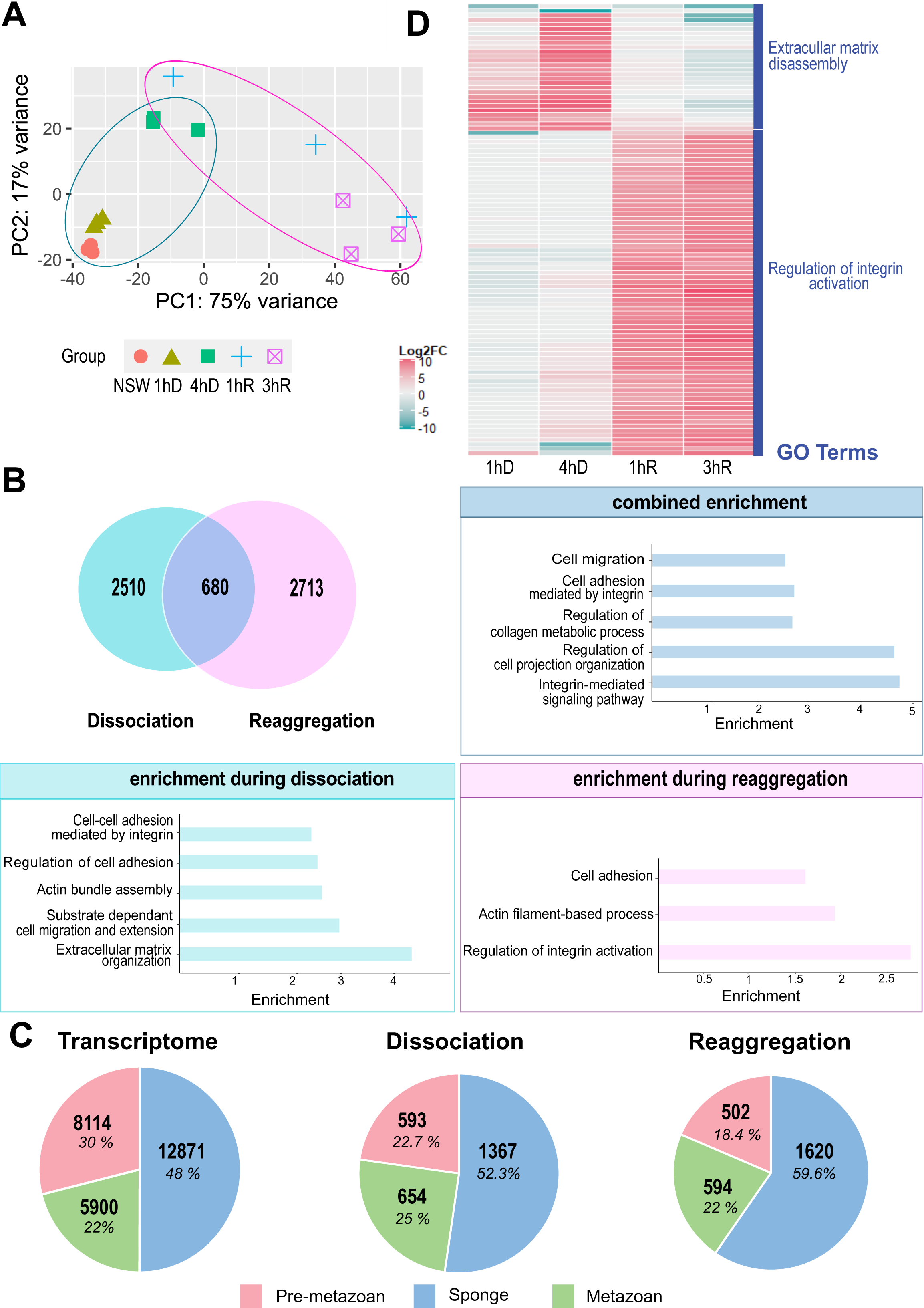
Gene expression changes in *O. lobularis* buds during dissociation and reaggregation. **(A)** Principal component analysis (PCA) plot of the 5 different conditions sequenced during dissociation and reaggregation. Dissociation time-points are circled in blue, those of reaggregation in pink. Replicates cluster well except for the samples of the first hour of reaggregation (1hR) which show larger dispersion, and which are located between time-points of four hours of dissociation (4hD) and three hours of reaggregation (3hR), indicating different reaggregation states of the sequenced buds. **(B)** A total of 5903 differentially expressed genes (DEGs) were found during dissociation and reaggregation with a log2 fold change (log2|FC|) of at least 1.5 (Supplementary Table S6). Pairwise comparisons were done as follows: for dissociation (1hD/CT, 4hD/CT); for reaggregation (1hR/4hD, 3hR/4hD); genes in common for both processes are shown as overlapping (680 genes). Gene ontology enrichment analysis of genes differentially expressed during dissociation shows that genes relevant for the process under study are enriched, including those involved in cell adhesion and its regulation, substrate dependent cell migration and actin bundle assembly were enriched (Supplementary Table S8). Enriched GO terms for shared DEGs of both conditions included cell migration, cell adhesion, regulation of collagen metabolic processes as well as integrin-mediated signaling pathway. During reaggregation, relevant GO terms associated with cell adhesion, regulation of integrin and actin filament-based process were enriched. (**C**) Phylostratigraphy analyses on the whole transcriptome and during dissociation and reaggregation provide the evolutionary age of differentially regulated genes during the dissociation and reaggregation process, as well as in the entire transcriptome of mixed stages. On the entire transcriptome 16.6% (5389) of the genes are *O. lobularis* specific (see Supplementary Table S7). During dissociation and reaggregation, there is a higher percentage of Sponge-specific DEGs as compared to the entire transcriptome. The percentage of metazoan DEGs is higher during dissociation than during reaggregation. Analysis was done after (Sogabe and Hatleberg, 2019). Species used are listed in Supplementary Table S7A. **(D)** Heatmap showing the hundred most differentially regulated genes during dissociation and reaggregation (1hD/CT, 4hD/CT, 1hR/4hD, 3hR/4hD). For genes upregulated during dissociation or reaggregation, we performed enrichment analysis for GO terms (with TopGO (Alexa, 2020)) (Supplementary Table S8B). Among the top enriched GO terms, we found regulation of extracellular matrix disassembly for dissociation and establishment and maintenance of an epithelium for the reaggregation.

#### Number and evolutionary emergence of genes involved in dissociation and reaggregation

We looked for differentially regulated genes between time-points by pair-wise comparisons using DESeq2 (Love et al., 2014). We found 5903 genes significantly differentially expressed (adjusted p-value < 0.05 and a log2|FC| of at least 1.5) between each time-point of dissociation and the NSW control and between each time-point of reaggregation and 4hD. The number of exclusively differentially expressed genes during dissociation (2510 genes) was comparable to the number of exclusively differentially expressed genes during reaggregation (2713 genes). 11.5% of differentially expressed genes (680 genes) were shared in both processes, dissociation and reaggregation (Figure 7B, see Supplementary Table S6A-C for the lists of differentially expressed genes of the processes under study).

To evaluate the proportion of Sponge-specific, Pre-metazoan or Metazoan genes differentially expressed during the dissociation-reaggregation process, we performed a phylostratigraphy analysis according to Sogabe et al. (2019). A phylostratigraphy analysis is based on sequence similarity with genes present in other organisms with a defined phylogenetic distance (Domazet-Lošo and Tautz, 2010). *O. lobularis* genes were classified in three categories of evolutionary emergence: (1) before or (2) after the divergence between choanoflagellates and metazoans (Pre-metazoan and Metazoan genes, respectively); and (3) after the divergence between the phylum Porifera and the other metazoan phyla (Sponge-specific genes) (Supplementary Table S7A, B). We first looked at the overall distribution of the phylogenetic category of conserved genes in the entire transcriptome and found that most of the genes (48%) were Sponge-specific and a slightly higher percentage of the remaining genes predated the metazoan emergence (30%), leaving 22% Metazoan genes (Figure 7C; Supplementary Table S7C). The proportion of genes in each phylogenetic category was slightly shifted in both dissociation and reaggregation: during both processes, we found slightly more Sponge-specific genes than present in the transcriptome (52.3% in dissociation and 59.6% in reaggregation, respectively). During both processes the percentage of Pre-metazoan genes decreased and the percentage of Metazoan genes slightly increased. We found a higher percentage of Metazoan genes (25% and 22% in dissociation and reaggregation, respectively) than Pre-metazoan genes (22.7% and 18.4% in dissociation and reaggregation, respectively) (Figure 7C; Supplementary Table S7C).

#### Identification of the molecular toolkit regulated during dissociation and reaggregation of a sponge

Among the 5903 differentially expressed genes, we next looked at enriched gene ontology (GO) terms associated with genes involved only in dissociation (2510 genes) or only in reaggregation (2713 genes), as well as those genes shared between both processes (680 genes) (Figure 7B; the complete list of enriched GO terms is available in Supplementary Table S8A). In order to investigate, if epithelial organization shares molecular components with bilaterians we focused on genes related to cell-cell contacts, cell adhesion and migration.

Genes involved in both, dissociation and reaggregation were enriched for GO terms related to cell-migration, control of cell projection and cell adhesion (including integrin-dependent mechanisms) and control of collagen metabolism (Figure 7B). For the significant transcripts associated with these GO biological processes, we identified the encoded genes by identifying orthologous groups with eggNOG (Huerta-Cepas et al., 2019), as well as by performing BlastP searches (Altschul et al., 1997) (Supplementary Table S9A). We observed that most of these genes encoded proteins involved in cell surface receptor tyrosine kinase or phosphatase activities.

During dissociation, mainly genes involved in regulation of cell adhesion, cell migration, actin bundle assembly were enriched, and even more genes involved in extracellular matrix organization. This latter category included in particular extracellular matrix-structuring proteins such as Collagens, Microfibril-Associated Glycoprotein (MAGP), Laminin or Papilin and Integrin receptors (Supplementary Table S9A).

Genes involved in reaggregation were enriched for GO terms related to cell adhesion, actin filament-based processes and in the regulation of integrin complex activation (Figure 7B). The latter GO category included genes such as Protocadherin Fat 4, Cadherin 23-like or Afadin (Supplementary Table S9A).

We finally looked at the 100 most differentially expressed genes over all four key steps of dissociation-reaggregation (Figure 7D). We found different sets of genes specific either for dissociation or for reaggregation. We used those genes for GO-term enrichment analysis (Supplementary Table S8B). During the dissociation process, in agreement with our previous results, the most differentially expressed genes were mainly associated with the GO category “Extracellular matrix disassembly”. For the reaggregation process, we found GO-terms related to “Regulation of integrin activation”. In addition, we noticed that transcripts belonging to two orthologous groups of C-type lectins (Supplementary Table S9B) were strongly increased during the reaggregation process.

In summary, we found biological processes related to cellular adhesion, cell migration, integrin signaling and actin assembly as well as processes important for organizing the extracellular matrix enriched in the differentially expressed genes during the *O. lobularis* dissociation-reaggregation processes.

### Gene expression dynamics during dissociation and reaggregation of *Oscarella lobularis* buds

To identify genes with a similar temporal expression profile, we performed Mfuzz clustering (Futschik and Carlisle, 2005) on standardized transcript per million read counts (TPM) of all expressed genes. In this analysis, the expression profile over time, rather than the degree of differential expression is decisive, allowing to monitor the overall expression dynamics of genes during dissociation-reaggregation and to identify co-regulated gene groups. Mfuzz cluster profiles are shown in Figure 8A (Supplementary Table S10 for the list of the core cluster genes). We could discriminate different types of expression profiles in our dataset: those with a decrease in expression during dissociation and upregulation of genes during reaggregation (clusters 12, 15); those with decreasing expression during the time-course (clusters 3, 8, 5, 6, 10); those with a strong increase of expression during reaggregation (clusters 1, 11, 13); finally, those with a peak in expression either during late dissociation or early reaggregation (clusters 7, 2, 9, 4, 14). Cluster cores, defined by a membership value α > 0.7, contained between 43 and 654 genes.

**Figure 8:**
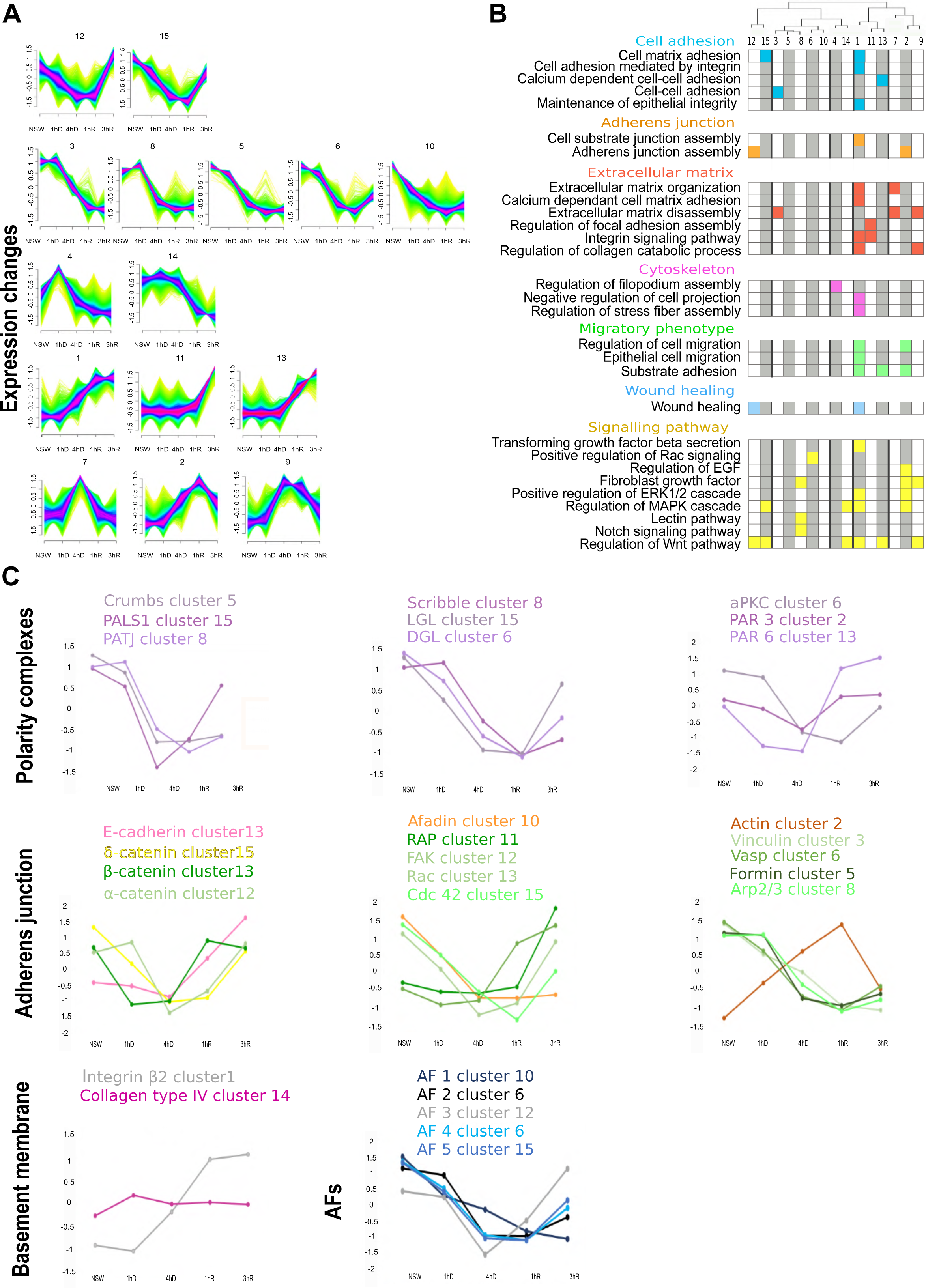
Gene expression dynamics during dissociation and reaggregation. (**A**) Fuzzy clustering (Futschik and Carlisle, 2005) of temporal expression profiles during dissociation and reaggregation. We chose 15 clusters to represent our data, determined by hierarchical clustering of normalized read counts of the five conditions using hclust in R (Supplementary Figure S7). We found subgroups of profiles with a decrease in expression until 4hR (clusters 12,15), early peaks and a flat curve during reaggregation (clusters 3,8,5,6,10), peaks at 1hR (clusters 4,14), decreasing expression during dissociation and increasing expression in reaggregation (clusters 1,11,13), strong ascending trends (clusters 10,5) and a peak at 4hD (clusters 7, 2, 9). Cluster cores have a membership value > 0.7 (pink) (Supplementary Table S10). Other membership values are shown as follows: membership value between 0.5 and 0.7 (blue), membership value < 0.5 (green). **(B)** Selected GO terms corresponding to Biological Processes found to be enriched for each core cluster as determined by topGO (Alexa, 2020)(Supplementary Table S8C). Colored boxes indicate significant enrichment in a given term (adj p-avlue <0.05). The dendrogram at the top was calculated by hierarchical clustering using hclust with Euclidian distance of the normalized read counts of genes qualifying as cluster core (membership value >0.7). **(C)** Expression profile of genes involved in epithelia organization and cell adhesion in bilaterians (see also Figure 1A). Expression of genes of the three polarity complexes decrease during dissociation and increase upon reaggregation. The expression of the gene coding for the basement membrane collagen was rather stable during the process, while the expression of integrin β2 increased at 4hD and became stable at 1hR. Genes involved in adherens junctions globally show a decrease in expression during dissociation and an increase around 1hR. Expression of the cytoskeleton component actin increased upon dissociation and decreased when reaggregation started. Finally, four of the Aggregation Factors (AFs) candidates (AF 1, 3, 4, 5) were downregulated during dissociation and upregulated during reaggregation (in particular AF3). In contrast, AF2 was downregulated during the entire process. NSW stands for natural condition which is the control condition, 1hD and 4hD correspond to 1 hour and 4 hours of dissociation (in CMFSW) respectively, 1hR and 3hR stand for 1 hour and 3 hours or reaggregation (in NSW) respectively. See Supplementary Table S11 for Standardized TPM and log2 Fold Change values.

We performed GO enrichment of core Mfuzz clusters using TopGO (Alexa, 2020) (Figure 8B; complete list of GO terms are available in Supplementary Table S8C). During both, the dissociation and reaggregation process, a large set of terms related to locomotion, cell migration, wound healing, adherens junction regulation, EGF-receptor signaling as well as MAPK signaling were enriched in different clusters: during dissociation we observed a decrease of transcripts related to lectins and the Notch pathway (Cluster 8), FGF signaling (Cluster 8) and an increase of transcripts related to filopodium assembly (Cluster 4), regulation of focal adhesion assembly (Cluster 11), TGFβ signaling (Cluster 1), as well as EGF and FGF signaling (Cluster 2,9). During reaggregation, genes belonging to enriched GO terms related to cell adhesion (involving Calcium-dependent mechanisms) (Cluster 13) were increased, and genes related to the regulation of extracellular matrix disassembly (Cluster 7, 9) and filopodium assembly (Cluster 4) were decreased.

In addition, we searched in clusters with common expression profiles for transcripts with conserved domains related to cadherin, integrin and immunoglobulins as major domains involved in the Cell Adhesion Molecules (CAM) (Harjunpää et al., 2019) (Supplementary Figure S4). According to the phylostratigraphy analysis, more than 30% of these transcripts were Sponge-specific (Supplementary Table S7D). Furthermore, these specific transcripts showed interesting profiles with decreased expression at 4hD and an increase during reaggregation.

We looked more specifically at the expression profiles of genes involved in epithelial organization in bilaterians, including genes encoding proteins involved in cell polarity, basement membrane, as well as adherens junctions (Figure 8C; standardized TPM values are provided in Supplementary Table S11A). The 9 genes that are part of the three polarity complexes Crumbs, Scribble and Par had similar expression patterns: they decreased during dissociation and increased again during reaggregation (Figure 8C). Par complex genes showed slightly different dynamics and recovered earlier in reaggregation to initial levels. Expression profiles of genes involved in forming adherens junctions followed a similar trend, with downregulation at early stages followed by increasing expression levels during the reaggregation process, though the variability of observed profiles was higher for this gene group. Notably, Actin showed a reverse trend to all adherens junction genes, with increasing expression levels during dissociation and a drop in expression levels between 1h and 3h of reaggregation (Figure 8C). The two genes coding for the basement membrane components Integrin β2 and type IV Collagen showed differential expression dynamics: type IV Collagen was mostly stable during the time-course with only a slight increase in expression at 1hD, suggesting that loss of protein immunostaining we observed (Figure 3) was likely controlled on protein level. In contrast, the expression of Integrin β2 was induced during late dissociation and became stable at 1hR.

Finally, we looked at Aggregation Factors (AFs) that are thought to be key players of cell adhesion processes in sponges (Harwood and Coates, 2004). The five putative Aggregation Factors (belonging to AF-Group3 according to the analyses of (Grice et al., 2017)), were recovered in clusters 10, 6, 12, 15 (Figure 8C; TPM standardized values are shown in Supplementary Table S11B; AF domain composition is shown in Supplementary Figure S5 and Supplementary Table S12). Nearly all AFs showed similar expression dynamics, with a decrease during dissociation and an increase in expression during reaggregation, with one AF being the exception that did not recover its transcript levels by 3hR (AF 1 in cluster 10).

In conclusion, we found an enrichment of terms related to cellular adhesion, locomotion, adherens junction, as well as cytoskeleton organization in several profile clusters. Moreover, several proteins known to be required for epithelial structures or epithelial organization in bilaterians, as well as the Sponge-specific aggregation factors were differentially regulated during the dissociation and reaggregation process.

## Discussion

### A new transcriptomic resource targeted to study ancestral epithelial actors

In this study, we present an improved version of the transcriptome of *Oscarella lobularis* (Schenkelaars et al., 2015). We used a mix of stages, which included an adult in reproduction which contained pre-larvae and embryos, as well as buds under different environmental conditions (calcium-magnesium deprivation or not) and combined different methods of sequencing (454 + Illumina). This new transcriptomic resource is therefore a valuable addition to the few (2 sequenced species out of 126 described homoscleromorph species) transcriptomic and genomic resources currently available for the homoscleromorph sponge class (Ereskovsky et al., 2017; Nichols et al., 2012; Riesgo et al., 2014, reviewed in Renard et al., 2018). The *de novo* assembled transcriptome of *O. lobularis* revealed a high degree of completeness (82% BUSCO score) and substantial similarity to other sponges and metazoans (56.6% of orthologs with the closest sponge species). It also contained a high percentage (72%) of conserved domain arrangements as determined by DOGMA. This is well within the range for other non-standard model organisms and other sponges. *A. queenslandica* has a conserved domain arrangement of 81.65%, *A. vastus* of 60.81% and this score is 75.65% for the closely related *O. pearsei*.

Nonetheless, it is probable that the *O. lobularis* transcriptome presented here is incomplete, best reflected in the total number of contigs and the percentage of conserved genes with other sponges and metazoan species. For example, it is currently estimated based on near-complete sponge genomes that sponges have a large gene repertoire and gene numbers range between ∼30000 and ∼40000 (Ereskovsky et al., 2017, Kenny et al., 2020). The closest neighbor of *O. lobularis*, *O. pearsei* has about 29000 predicted genes (Ereskovsky et al. 2017). Based on the analysis of our transcriptome, we only find 56% shared genes between these two closely related species. The *O. lobularis* transcriptome shows a relatively high number of Species-specific genes (> 40%) compared to other Porifera (Riesgo et al., 2014). While we tried to reduce redundancy in our assembly by clustering similar contigs using CD-HIT, remaining redundant contigs, or only partially assembled transcripts could obscure the annotation process and fail to assign orthologs between species. It is therefore desirable to aim for a chromosome-level assembly of this sponge. Due to the relatively high number of repetitive elements this genome seems to harbor, this could be achieved by combining long-read sequencing with partial genome sequence data already available, partial genome sequence data (Belahbib et al., 2018). This resource presented here should then help in predicting genes with high accuracy. Furthermore, for sponge-specific transcripts, it is desirable to study those with interesting expression patterns derived from our experimental setup functionally.

Beside the *de novo* transcriptome provided here, this study is the first transcriptomic analysis designed to identify genes involved in cell dissociation-reaggregation in a sponge, despite the long-standing interest for this process in sponges (Wilson, 1907). This particular experimental context is especially suited to explore molecular mechanisms involved in epithelial morphogenetic processes in sponge, to determine, whether these processes are conserved between sponges and bilaterians and thus, to study the evolutionary origin of epithelium formation.

### Calcium as key regulator for epithelium morphogenesis

Cell dissociation in the homoscleromorph sponge *Oscarella lobularis* was induced by incubating biological samples in Calcium and Magnesium Free Sea Water. While this type of experiment was already performed in numerous calcareous sponges and demosponges (Curtis, 1962, 1970; Galtsoff, 1925; Humphreys, 1963; Huxley, 1921; Lavrov and Kosevich, 2014, 2016; Leith, 1979; Maclennan and Dodd, 1967; Simpson, 1984; Wilson, 1907), this study is the first one i) performed on a homoscleromorph, ii) with such a level of precision regarding the cellular description, and iii) using transcriptomic approaches to decipher molecular mechanisms involved during these processes. This taxonomic choice is key since homoscleromorph sponges are the only ones to possess epithelial layers fitting the three criteria found in Cnidaria and Bilateria (namely cell-polarity, cell junction plus a basement membrane). Consistent with the role of calcium deprivation described in other metazoans, cell-cell and cell-matrix interactions and epithelial integrity are affected in *O. lobularis* in the same way as in other animals resulting in the loss of epithelial characteristics.

Under calcium-magnesium deprived conditions, the choanocytes, forming the choanoderm epithelial layer 1) change their cell shape (from conical to round shape) and lose their basal-apical polarity; 2) reorganize their actin cytoskeleton (loss of apical ring of actin microvilli and increase of actin-based protrusions at the basal pole); 3) lose their cell-cell and cell-matrix contacts; and 4) start to migrate (towards the choanocyte chamber internal cavity or in the mesohyl).

Both the loss of tight cell-cell and cell-matrix contacts and the reorganization of the actin cytoskeleton are highly reminiscent of the phenotype resulting from calcium and magnesium deprivation in mammalian and other bilaterian epithelial cells. It is well known that, in bilaterians, cell contacts are calcium-dependent and removing bivalent cations from the medium induces junction disassembly and detachment from the basal lamina (Brodskiy and Zartman, 2018; Brown and Davis, 2002; D’Souza et al., 2019; Giannone et al., 2004; Ko et al., 2001; Naik and Naik, 2003; O’Keefe et al., 1987).

In addition, this process is reversible, as is also reported for bilaterians: incubation in natural sea water induces cell-reaggregation and re-epithelialization with a complete restoration of cell polarity, cell-cell and cell-matrix contacts. Such a plasticity of epithelial tissues is essential since epithelial cell movements are required during major morphogenetic processes such as, for example, wound healing and regeneration, tissue renewal, or gastrulation (Donati and Watt, 2015; Kim et al., 2017; Nieto, 2013; Plygawko et al., 2020). In bilaterians, calcium signaling is commonly and naturally involved in the regulation of epithelial tissue properties (motility, differentiation, cilia beating) during developmental processes (Brodskiy and Zartman, 2018). Interestingly even if calcium-signaling is an ancient feature of Eukaryotes, the emergence of a complex epithelial level of organization coincides with the multiplication and diversification of calcium-signaling proteins involved in cell-cell contacts in animals (Marchadier et al., 2016). The calcium depletion implemented in this study in a so-called « epithelio-sponge » (Ereskovsky et al., 2013) provides an invaluable opportunity to study the evolution of calcium-dependent molecular mechanisms involved in epithelial morphogenesis during animal evolution.

### An ancestral metazoan molecular toolkit involved in epithelium organization and morphogenesis

In bilaterians, the regulation of epithelial features involves an intimate interplay between adhesion proteins, polarity protein complexes and regulators of the actin cytoskeleton. The current knowledge on sponge epithelia suggests that they present similarities with those of bilaterians: chemical and mechanical properties (Elliott and Leys, 2007; Leys et al., 2019), histological likeness (Ereskovsky, 2010; Leys and Hill, 2012; Leys and Riesgo, 2012; Leys et al., 2009), supposed involvement of Wnt signaling in their regulation (Adamska, 2015; Adamska et al., 2007, 2010; Lapébie et al., 2009; Schenkelaars et al., 2015, 2016; Windsor and Leys, 2010; Windsor Reid et al., 2018) and shared epithelial genes and proteins (Belahbib et al., 2018; Fahey and Degnan, 2010; Green et al., 2020; Miller et al., 2018; Mitchell and Nichols, 2019; Nichols et al., 2012; Riesgo et al., 2014; Schippers et al., 2018). Nevertheless, the scarcity of available data prevents a comprehensive and global vision of genes involved either in the composition or in the regulation of sponge epithelia. The homology and functional conservation of proteins involved thereby remains an open question. Our presented molecular data from transcriptomic analysis during an epithelial morphogenetic process brings important new insights on this key evolutionary question.

#### Crumbs as an ancestral regulator of epithelial features?

The protein Crumbs is a metazoan innovation (Assémat et al., 2008; Bazellières et al., 2018; Belahbib et al., 2018; Le Bivic, 2013; Renard et al., in press). In bilaterians, Crumbs is not only involved in the establishment and maintenance of cellular apico-basal polarity and the positioning and stability of adherens junctions, but is an essential player in the dynamics of actin remodeling by interacting with many actin binding partners (Bazellières et al., 2018). These partners can interact with the FERM or the PDZ-binding domains of Crumbs directly or indirectly. For example, Crumbs proteins can regulate actin dynamics by recruiting ARP2/3 or by regulating Cdc42 (Bazellières et al., 2018). In *O. lobularis,* orthologs shared with the bilaterian proteins involved in the Crumbs complex (namely Crumbs, Patj, Pals1) have been reported, and key functional motifs required for their functional interactions have been clearly identified (Belahbib et al., 2018). Here we evidence that the genes encoding Crumbs, Pals1 and Patj are downregulated during dissociation and upregulated during the reaggregation (only Crumbs is differentially expressed at 4hD) (Figure 9; Supplementary Table S11A for Log2 Fold Change values). Both, our previous *in silico* analysis (Belahbib et al., 2018) and their very similar expression dynamics during dissociation and reaggregation, are consistent with the conservation of their joint functional involvement in epithelium integrity, as has been shown in other animals (Assémat et al., 2008). Because these genes, as well as an epithelium in general, are specific to metazoans, it suggests that Crumbs and its partners have an ancestral role in epithelial organization in Metazoa.

**Figure 9:**
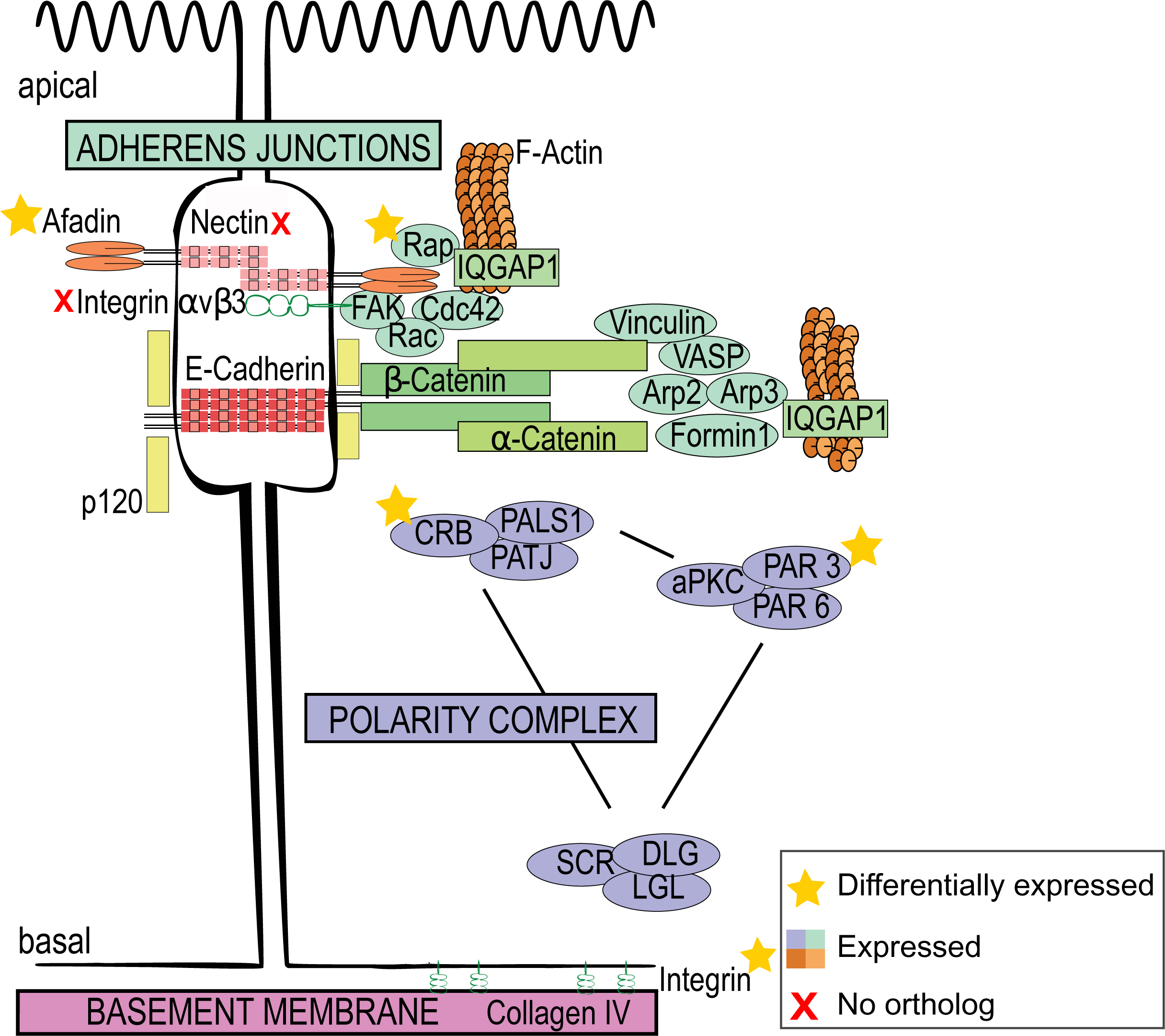
Gene expression of epithelial genes during the dissociation-reaggregation in *Oscarella lobularis*. Schematic representation of a bilaterian epithelial cell (Figure 1A) with the expression of the sponge genes coding for proteins involved in epithelial adhesion and maintenance during the dissociation-reaggregation experiment. For the genes involved in the adherens junctions, Afadin and Rap are differentially expressed at 4hD and 3hR, respectively. No orthologs were found for the Nectin and the Integrin αvβ3. In the polarity complex, Crumbs and Par3 are differentially expressed at 4hD. For the genes involved in the basement membrane, Integrin β2 is differentially expressed at 4hD. See Supplementary Table S11 for Standardized TPM and Log2FC values.

Similarly, the genes encoding the proteins involved in the two other bilaterian polarity complexes, the Scribble and Par complexes, are also downregulated while epithelium characteristics are lost; and then upregulated during the re-acquisition of epithelial features. The fact that i) all these genes (except Dlg) are metazoan innovations (Renard et al., in press), ii) their protein domain structures are highly conserved (Belahbib et al., 2018) and iii) their expression dynamics during the epithelial morphogenetic process studied here advocate for their ancestral joint involvement in metazoan epithelia. This also suggests that the more ancient Dlg gene (supposed Holozoan origin, (de Mendoza et al., 2010)) was co-opted secondarily for epithelial functions in the Last Metazoan Common Ancestor (LMCA). Further expression, localization and functional studies will help decipher the exact role of these genes in sponges, and thereby help resolve their respective ancestral roles.

#### Cadhesome, integrin-based “adhesomes” and cytoskeleton

In bilaterians, the crucial cross talk between cell-cell and cell-matrix adhesion sites for epithelium stability and plasticity is well described. Cadherin- and integrin-dependent adhesion and signaling functions intersect and interact. Changes in adhesion/force transmission at one site may affect membrane trafficking, cytoskeletal association, avidity or binding affinity (Bays and DeMali, 2017; DeMali et al., 2014; Weber et al., 2011). Integrins and cadherins are both transmembrane adhesion receptors. They have many signaling effector molecules in common and are linked to common scaffolding and cytoskeletal elements. Both adhesion molecules are essential for the formation of cell-cell and cell-matrix contacts through adherens junctions (AJs) and focal adhesions (FAs), respectively. It is therefore not surprising that during epithelium disassembly-reassembly, the genes coding for integrins are differentially expressed. The expression of most of the genes involved in both, cadherin and integrin adhesion complexes, are downregulated during dissociation and upregulated during reaggregation. These proteins are therefore likely to play important roles in epithelial organization in sponges as they do in bilaterian species. In addition, our results show that many genes encoding extracellular matrix proteins displayed differential expression, such as Glycoproteins, Laminins or Disintegrin and Metalloproteases (ADAMs), which are integrin ligands (Bridges and Bowditch, 2005; Giebeler and Zigrino, 2016). These results suggest that, like in bilaterians, the Extracellular Matrix (ECM) of *O. lobularis* plays an important role as substrate for migration, assembly, and regulation of the epithelial cells (Rozario and DeSimone, 2010; Scott et al., 2019).

Together with previous genomic surveys (Belahbib et al., 2018; Nichols et al., 2006) and recent biochemical studies (Miller et al., 2018; Mitchell and Nichols, 2019; Schippers et al., 2018), our results strengthen the hypothesis that cadherin and integrin-based mechanisms are ancient features of metazoans, as seems their ancestral roles in cell and tissue cohesion in the LMCA. Interestingly, the E-cadherin gene is barely changing expression levels during cell dissociation while it shows an increase during reaggregation and expression of the afadin gene shows opposite regulation. This unexpected finding according to previous results (Baj et al., 2020; Kaszak et al., 2020) questions the relative contribution of Cadherin and Afadin in the establishment and the maintenance of adherent junctions in *O. lobularis*. It cannot be excluded that the expression of these genes is regulated at protein level. Immunolocalization of their encoded proteins will shed light on this issue.

Beyond what is known about other metazoans, different genes may be involved in adhesion in sponges. During dissociation-reaggregation, some genes encoding other proteins involved in adhesion show a differential expression: this is the case for genes encoding Fat 4 proto-cadherin, non-classical Cadherins 23 and C-type lectins. For example, the increase of C-type lectin domain containing genes during reaggregation is in full agreement with its reported functions in bilaterians: i) as suppressor of Epithelial-Mesenchymal Transition (EMT) (Rodriguez-Teja et al., 2015); ii) as being involved in adhesion in budding tunicates (Matsumoto et al., 2001); iii) as necessary for cell-aggregation in choanoflagellates (formation of rosettes) (Levin et al., 2014); and iv) as being involved in cell-reaggregation in the sponge *Aphrocallistes vastus* (Hexactinellida) (Gundacker et al., 2001; Muller et al., 1984). In vertebrates, C-type lectin domain proteins are known to facilitate the Ca ^2+^-dependent cell-matrix and cell-cell interactions (Rodriguez-Teja et al., 2015). In the same way, Fat 4 and Cadherin 23 proteins were shown to regulate cell-cell adhesion and to be involved in dynamic changes leading to epithelial fluid-like attributes in bilaterians (Apostolopoulou and Ligon, 2012; Kumar et al., 2019).

#### Signaling pathways with conserved roles in the regulation of epithelial morphogenesis

The transition from unicellularity to multicellularity does not only rely on cell adhesion but also on the capability of neighbor cells to communicate and synchronize and/or to separate cellular tasks. The emergence of intercellular signaling provided cells with the means for cell-cell communication. Several signaling pathways play important roles in the regulation of metazoan morphogenetic processes: these include the canonical Wnt, the Hh (Hedgehog), the JAK/STAT (Janus kinase/signal transducer and activator of transcription), the Notch/Delta, the NRs (nuclear receptors), the RTK (receptor tyrosine kinase) and the TGF-β (transforming growth factor β) signaling pathways (Babonis and Martindale, 2017). Four of them (Wnt, NRs, Notch and TGF β) are considered present in sponges and represent innovations that probably emerged in the LMCA (Babonis and Martindale, 2017; Renard et al., 2018; Schenkelaars et al., 2019). During the dissociation-reaggregation processes, we noticed that members of several signaling pathways were differentially expressed. These included members of the Notch, TGFβ, NRs and Wnt pathways, as well as genes involved in EGF (epidermal growth factor) and the FGF (Fibroblast growth factor) signaling. A joint involvement of TGF-β, EGF, FGF, Wnt, Notch and integrin signaling may recall what occurs during epithelial-mesenchymal transition (EMT) (Gonzalez and Medici, 2014). Though the herein considered context is different, we cannot exclude common signaling mechanisms driving epithelial cell reprogramming.

Among these pathways, the canonical Wnt pathway (Wnt/β-catenin) plays a central role in tissue patterning (Pond et al., 2020). Indeed, the cross talk between the canonical Wnt pathway, actin-cytoskeleton and cell-cell/cell-matrix junctions is well known to modulate the stability and the plasticity of epithelia in bilaterians (Sedgwick and D’Souza-Schorey, 2016). Our data, together with other studies of Wnt signaling in sponges and other non-bilaterians strongly support the ancestral involvement of Wnt signaling in epithelium patterning: i) the only sponges and cnidarians missing Wnt genes also lack epithelia (Chang et al., 2015; Schenkelaars et al., 2017) ii) Wnt signaling disturbance by either pharmacological or siRNA approaches leads to phenotypes consistent with incorrect morphogenesis of epithelial layers (choanoderm and/or pinacoderm). However, the cellular mechanisms impacted are unknown (Lapébie et al., 2009; Windsor and Leys, 2010; Windsor Reid et al., 2018).

The change in the expression of genes of the Wnt pathway observed during the present dissociation-reaggregation experiments suggests possible functions in the regulation of cell motility, cell adhesion and tissue patterning, as has been shown for bilaterians (Pond et al., 2020; Rosenbauer et al., 2020; Steinhart and Angers, 2018).

### Sponge-specific genes involved in epithelium organization open new perspectives on epithelium evolution

Among Sponge-specific genes, we chose to focus on Aggregation Factors (AFs), because of the historical significance of these extracellular proteoglycans concerning sponge dissociation-reaggregation experiments (Henkart et al., 1973; Humphreys, 1963, 1965; Leith, 1979; Moscona, 1963, 1968; Müller and Zahn, 1973; Müller et al., 1978, 1985; Muller et al., 1976). AFs were shown to be involved not only in allorecognition but also in physical cell-cell bridges-like connections. They play key species-specific cell adhesion roles. A recent study questioned the presence of AFs in the Homoscleromorph class because of the too low sequence similarity with demosponge AF sequences and because of the lack of evidence of reaggregation properties in this sponge class (Grice et al., 2017). In this study we identified 5 genes in *O. lobularis* considered as possible AFs candidates (AFs of Group 3) by Grice and co-workers (Grice et al., 2017). Interestingly, the transcripts of the 5 AF candidates decreased during the dissociation process and 4 of them were upregulated during reaggregation. This expression is consistent to observations described for demosponge AFs. Though this finding argues that these genes encode *bona fide* aggregation factors in this homoscleromorph sponge, biochemical studies are needed to confirm this hypothesis.

Interestingly, these aggregation factors have been proposed to be able to interact with the integrin RGD motif and to activate integrin signaling (Fernàndez-Busquets et al., 1998; Harwood and Coates, 2004; Mitchell and Nichols, 2019; Wimmer et al., 1999). Therefore, models that will be proposed for the establishment of cell-cell contacts and adhesion in sponges will have to consider and integrate the involvement of AFs and their possible interactions with other adhesion receptors.

In addition, the search for genes harboring cadherin, integrin and immunoglobulin conserved domains (usually found in the Cell Adhesion Molecules (CAM) (Harjunpää et al., 2019)) in the different clusters of expression has highlighted the presence of these domains in numerous Sponge-specific transcripts (more than 30%). While some of those might be resulting from partial redundancy in our *de novo* assembled transcriptome, some could be potential new candidates to play a role in the epithelial cell adhesion because of their increase in expression during reaggregation. However, to test the validity of these candidates, additional in-depth analyses of their domain composition and functional analyses will be needed.

## Conclusions

The transcriptomic data collected during dynamic epithelial processes provide new clues for understanding the molecular elements of epithelization in *Oscarella lobularis*, and, more generally, for investigating the evolution of the mechanisms involved in epithelia formation in bilaterians. This study supports the hypothesis that epithelium organization in *O. lobularis* involves both, genes and processes homologous to bilaterians, as well as Sponge-specific ones. Biochemical and functional experiments will be needed to assess to what extent the complex signaling cascades and protein interactions described in bilaterian epithelia organization predate the cnidarian-bilaterian emergence.

## Methods

### Animal husbandry

Adult specimens of *Oscarella lobularis* (Schmidt 1862) were collected by SCUBA diving in the bay of Marseille (France) and kept in a thermostatic chamber at 17°C. Budding was induced by cutting adults and each fragment was placed in a well containing 8 ml Natural Sea Water (NSW) as described in (Rocher et al., 2020; Borchiellini et al., 2021). Free buds are transferred into Petri dishes containing NSW (renewed once a week) and maintained at 17°C (Figure 2).

### Cell staining

To monitor choanocytes, fluorescein labeled lectin PhaE (*Phaseolus vulgaris Erythroagglutinin*) (Vector lab) was used to specifically stain this cell type (1: 500; Rocher et al., 2020). Buds were incubated overnight (ON) at 17°C then rinsed three time in NSW.

### Cell dissociation and reaggregation experiments

To induce cell-dissociation, stage 3 buds (8-29 days, (Rocher et al., 2020)) from *O. lobularis* (n=3) were placed in 24-wells culture plates containing 2 mL of Calcium Magnesium Free Sea Water (CMFSW) at 17°C. Choanocyte chambers integrity was monitored (see next section) at 20 minutes, 1 hour, 2, 3, 4 hours. According to our observations (see result section) the pivotal time-points chosen for further analyses were 1 hour of dissociation (time-point hereafter referred to as 1hD) and 4 hours (4hD).

After 4 hours of dissociation in CMFSW, cell-reaggregation was triggered by putting back buds into 2 mL of natural sea water (NSW). The reaggregation process was monitored after 30 min, 1 hour, 2 hours, 3 hours, 5 hours, 6 hours and 24 hours. According to our observations the pivotal time-points chosen for further analyses were 1 hour (1hR), 3 hours (3hR) and 24 hours of reaggregation (24hR). For each time-point (except 24hR), one part of the treated and NSW control samples were collected and frozen (-80°C) for RNA-sequencing and one part preserved in fixative solutions for imaging (see next section and Figure 2).

### Microscopy observations

At each stage of the dissociation (1hD, 4hD) and reaggregation (1hR, 3hR, 24hR) processes treated and control buds were observed by fluorescent light and electronic microscopy (except for reaggregation process), focusing on the choanocyte chamber.

For Scanning Electron microscopy (SEM), samples were prepared using the NCMIR protocol for SBF-SEM (Deerinck et al., 2010). Imaging was carried out on a FEI Teneo VS running in low vacuum (30 Pa), at 2kV and using a backscattered electron detector. Acquisition pixel size was 20x20x60 nm.

For confocal observations, samples were fixed overnight in 3% paraformaldehyde (PFA) in PBS at 4°C, rinsed twice and staining with DAPI (1:500) (Thermo Fisher Scientific) and 1:1000 (Alexa fluor 647 coupled-phalloidin (Santa Cruz Biotechnology) and mounted in Prolong Diamond antifading mounting medium (Thermo Fisher Scientific) as described in Rocher et al., 2020 and Borchiellini et al., 2021. Confocal acquisition was done with a Zeiss LSM 880 with a 63x Oil objective and images were processed using FIJI (Schindelin et al., 2012). All experiments were repeated at least 3 times.

### Type IV Collagen antibody production and validation

Antibodies against the type IV Collagen of *O. lobularis* were obtained by immunizing rabbits using the speedy protocol from Eurogentec (Seraing, Belgium) with a synthetic peptide (QTISDPGEEDPPVSKC) coupled to the KLH (keyhole limpet hemacyanin) carrier. This peptide is present in the C-terminal NCI domain of *O. lobularis* type IV Collagen and was used to purify antibodies from immunized rabbit sera.

To validate this antibody, we performed competition assays. During these assays, the antibodies against type IV Collagen were incubated ON at 4°C with either the immunizing peptide, or an unrelated peptide (CSTVSVAQTGLKGGGI, from the Crumbs protein, negative control) or only PBS buffer (positive control). The molar ratio between antibodies and peptides was 1 mole for 25 moles, respectively, and antibodies were used at 1:200 (2.6 μg/mL). After centrifugation (21000g, 4 hours, 4°C) to get rid of potential antibody-peptide complexes, supernatants were used to perform the immunostaining protocol (see next section). Incubation of antibodies with the immunizing peptide yielded to the loss of immunostaining unlike incubation with irrelevant peptide antibodies α-type IV Collagen (Supplementary Figure S6).

### Acetylated tubulin and type IV Collagen immunostaining

For immunofluorescence, buds are fixed in 3% PFA in PBS ON at 4°C. To monitor the epithelial features of the choanoderm, the type IV Collagen was immunolocalized to observe cell-matrix adhesion, acetylated tubulin (Sigma T6793) was immunolocalized to observe alteration of cell polarity and Alexa fluor-647 coupled-phalloidin (Santa Cruz Biotechnology) was used to follow actin-rich adhesive cell-cell junctions and cytoskeleton remodeling. Immunofluorescence was performed as described in Rocher et al. (2020) and additional technical details are available in Borchiellini et al. 2021.

### Statistical analyses of confocal acquisition

Perturbations of the choanoderm layer in CMFSW-treated buds compared to the controls were estimated according to 2 criteria (1) after 1 hour of dissociation (1hD) by the presence or absence of microvilli on choanocytes (2) after 4 hours of dissociation (4hD) and 24 hours of reaggregation by the integrity or not of the choanocyte chambers (cell-cell and cell-matrix adhesion, presence/absence of central cavity). The experiments were at least repeated 3 times for each time-points with at least 3 buds. The integrity of the choanoderm, according to the previously defined criteria, was observed by counting the choanocyte chambers on two distinct part of the buds (1hD: 3x4 buds, 178 choanocyte chambers counted; 4hD: 6x 2/3 buds; 216 choanocyte chambers counted; 24hR: 3x3 buds; 94 choanocyte chambers counted) (Supplementary Table S1). A Wilcoxon test was preformed using R (RStudio Team, 2015).

### RNA-sequencing

Biological triplicates of NSW control buds, 1hD, 4hD, 1hR and 3hR were collected and RNA was directly extracted with the RNAeasy Mini Kit (Quiagen). For each replicate, 40 clonal stage 3 buds per condition were pooled to perform RNA extraction. Sequencing was realized by the Transcriptomic Genomic platform of Marseille Luminy (TGML), using the Illumina Truseq stranded mRNA kit for library preparation and 75 nt single-end sequencing with the Illumina NextSeq 500 instrument, and 400 million reads per run. Raw sequencing data were submitted to the Submission Raw Archives (SRA) (SRR12538867, SRR12538868, SRR12538869, SRR12538870, SRR12538871, SRR12538872, SRR12538873, SRR12538874, SRR12538875, SRR12538876, SRR12538877, SRR12538878, SRR12538879, SRR12538880, SRR12538881, SRR12538882).

### *De novo* assembly of the *O. lobularis* transcriptome

The *de novo* transcriptome assembly strategy is shown in Figure 5. In brief, 454 sequencing data from the adult in sexual reproduction with embryos and larvae (Schenkelaars et al., 2015) and illumine sequencing data from *O. lobularis* buds were first assembled individually and then combined to obtained a transcriptome containing transcripts from all life stages. The 454 dataset of adult *O. lobularis* was assembled with the Staden package and GAP4 (Bonfield et al., 1995) (v1.7.0) by Eurofins. Illumina reads were first quality filtered using FASTQC (v 0.11.8, available at http://www.bioinformatics.babraham.ac.uk/projects/fastqc/) and trimmed using TrimGalore (v0.6.4, available at https://github.com/FelixKrueger/TrimGalore). Trimmed sequences were *de novo* assembled using Trinity (v2.9.1, (Grabherr et al., 2011)). The two datasets were concatenated and clustered using CD-HIT EST (v4.8.1, (Li and Godzik, 2006)), whereby all sequences with 80% sequence identity were clustered together and the longest transcript was kept. We also used GMAP (v2019.12.01, (Wu and Watanabe, 2005)) to remove all shorter sequences that were 100% identical to a longest sequence. Vector contamination were removed with Vecscreen ((Hancock and Bishop, 2014), available at https://www.ncbi.nlm.nih.gov/tools/vecscreen/) using default parameters: this tool performs a BLAST+ (BlastN (Camacho et al., 2009)) search against the UniVec database. To remove sequences that might have contaminated our biological specimen, we performed BlastN searches against a custom database composed of Archaea, Bacteria, Viridiplantae, Fungi and Dinoflagellata (e-value 10^-6^ and a percentage of identity of 80%). Protein coding sequences were predicted using Transdecoder (v5.5.0, available at https://github.com/TransDecoder). By default, Transdecoder will identify ORFs that are at least 100 amino acids long. We cross-validated Transdecoder results with results from HMMscan ((Eddy, 1998), available at http://hmmer.org/) against the PFAM database ((El-Gebali et al., 2019), available at http://pfam.xfam.org/) for predicting the most likely coding regions. The completeness of the transcriptome was evaluated by BUSCO (v4.1.2, (Seppey et al., 2019). BUSCO scores were calculated based on the BUSCO Metazoan gene set. In addition we tested the completeness of conserved domain arrangements with DOGMA ((Dohmen et al., 2016) ,https://domainworld-services.uni-muenster.de/dogma/) (Supplementary Table S4 and S5).

### *Gene functional annotation of the* de novo *assembled* O. lobularis *transcriptome*

We annotated gene functions by information transfer from orthologous sequences. Orthologous groups were identified with eggNOG mapper (v5.0, (Huerta-Cepas et al., 2019)). The percentage of orthologous groups that *O. lobularis* has in common with other metazoans species was inferred by Orthofinder searches (v2.2.7, (Emms and Kelly, 2019)) against a set of selected species: *Homo sapiens* (GRCh38.pep from ENSEMBL), *Drosophila melanogaster* (BDGP6.28.pep from ENSEMBL), *Nemastostella vectensis* (ASM20922v1.pep from ENSEMBL), *Trichoplax adhaerens* (ASM15027v1.pep from ENSEMBL), *Sycon ciliatum* (SCIL_T-PEP_130802 from COMPAGEN), *Oscarella pearsei* (OCAR_T-PEP_130911 from COMPAGEN) *Amphimedon queenslandica* (Aqu1.pep from ENSEMBL) and *Aphrocallistes vastus* (AV4_1_Trinity from ERA archive of Alberta university) and *Salpingoeca rosetta* (UP000007799 from Uniprot). Functional annotation with GO terms was done using eggNOG and InterProScan (v5.40, (Jones et al., 2014)). Identification of conserved domains was done using CD-search (CDD-database version 3.18, (Marchler-Bauer et al., 2011)).

### *Identification of Aggregation Factors from the* Oscarella lobularis *transcriptome*

*Oscarella lobularis* aggregation factors were identified using BlastP (e-value=10) with *Oscarella carmela* aggregation factor candidates from group 3 identified in the study of Grice (Grice et al., 2017) as queries. Protein domain predictions were done using Interproscan online with default parameters for each sequence (Supplementary Figure S5 and Supplementary Table S12).

### Differential and time-series analysis of RNA-seq data

Illumina sequences were mapped using Kallisto (v0.46.2, (Bray et al., 2016)) and Tximport (Soneson et al., 2015) to resolve ambiguities in read mapping. Detection of differentially expressed genes was performed with DESeq2 (Love et al., 2014). Lowly expressed genes, defined by having less than 10 read counts across all samples, were excluded. Genes with padj < 0.05 and log2 fold change (log2|FC|) of at least 1.5 were defined as differentially expressed. Pairwise comparisons were done as follows: dissociation time-points 1hD and 4hD were compared to NSW control and reaggregation time-points 1hR and 3hR were compared to the last time-point of dissociation, 4hD.

Soft clustering of time-course data was performed in R with Mfuzz (Futschik and Carlisle, 2005), using the transcripts per million (TPM) count table (Supplementary Table S13) after a standardization. Hierarchical clustering, using the hclust function in R (Supplementary Figure S7), was performed to evaluate the number of clusters parameter needed as input for the Mfuzz clustering. By inspecting the dendrogram, 15 clusters showed to well discriminated the data. The fuzzifier value m prevents clustering of random data, is the second parameter needed for the algorithm, was calculated with the function mestimate. This value has been estimated at 1.99. Mfuzz clustering was repeated 10 times to test the clusters stability by calculating the Jaccard index to evaluate the similarity of the core cluster between each run (Supplementary Table S14). Mfuzz core clusters were restricted to genes with a membership value (α) greater or equal to 0.7; to visualize the relationship between clusters (Figure 8), average expression profiles of member genes were calculated as the average standard-normal expression and then were clustered in R using Euclidean distance and complete linkage.

### Enrichment analysis

Enrichment analysis was performed with the TopGO package in R (available at https://bioconductor.org/packages/release/bioc/html/topGO.html) using the weight 01 algorithm, a classic Fisher’s exact test on the Biological Process GO category with the annotated transcriptome as background. Enrichment of Conserved Domains was done with the clusterProfileR package on custom made Conserved Domain IDs (Yu et al., 2012) using a Fisher’s exact test. Full results and gene lists are available in the (Supplementary Table S8).

### Phylostratigraphy analysis of the O. lobularis transcriptome and differentially expressed genes during dissociation and reaggregation

The evolutionary emergence of genes of the transcriptome and those regulated during the dissociation and reaggregation of *O. lobularis*, was estimated using the method proposed by (Sogabe et al 2019) with a custom database containing 27 transcriptomes and proteomes (Supplementary Table S7). Identification of orthologous groups with eggNOG (Huerta-Cepas et al., 2019) and BlastP (Camacho et al., 2009) searches was performed for every coding sequence of *O. lobularis* against this custom database, and its evolutionary age was inferred based on the oldest Blast hit relative to the predetermined phylogenetic classification of all 27 species (Supplementary Table S9).

### Data availability

This Transcriptome Shotgun Assembly project has been deposited at DDBJ/EMBL/GenBank under the Bioproject PRJNA659410. The version described in this paper is the first version publicly available. Supplementary Table S2 contains all the information available for the transcriptome. Translated version of the transcriptome is available in Supplementary File S1.

## Declarations

### Ethics approval and consent to participate

’Not applicable’ for that section.

### Consent for publication

’Not applicable’ for that section.

### Availability of data and materials

All data generated or analysed during this study are included in this published article and its supplementary information files. Raw files of the sequencing and transcriptome assembly has been deposited at DDBJ/EMBL/GenBank under the Bioproject PRJNA659410.

## Authors’ contributions

AV performed the experiments, participated in transcriptome assembly and analysis, and writing of the manuscript. MMP participated in transcriptome assembly and analysis, in writing of the manuscript. FM participated in transcriptome and time-series analysis helped edit the manuscript. HD was involved in validation of the antibody. NB participated in electron microscopy (SEM 3D) imaging and helped edit the manuscript. DMH participated in validation of the antibody and manuscript editing. CR provided technical support and participated in manuscript editing. ER was involved in project design, data analysis and manuscript writing. ALB was involved in project design, data analysis and manuscript writing. BHH was involved in bioinformatic project design, data analysis and manuscript writing. CB was involved in project design, data analysis and manuscript writing. All authors read and approved the final manuscript.

## Acknowledgements

We would like to thank the OEB Team of the IMBE lab for the financial support concerning the RNA sequencing; the IMBE molecular biology service, the IMBE morphology service, the diving service of OSU Institut Pytheas and divers from the IMBE lab (Christian Marschal, Sandrine Chenesseau) for sampling and their time and advices. We are grateful to the Genotoul bioinformatics platform Toulouse Midi-Pyrenees (Bioinfo Genotoul) and High-Performance Computing (HPC) cluster of the Institut Pytheas (OSU) for providing computing resources. A special thanks to all members of the Habermann Team (IBDM) for all their precious advices and technical supports.

## Authors’ information

’Not applicable’ for that section.

## Footnotes

’Not applicable’ for that section.

